# A domestic plant differs from its wild relative along multiple axes of within-plant trait variability and diversity

**DOI:** 10.1101/2020.11.14.382788

**Authors:** Moria L. Robinson, Anthony L. Schilmiller, William C. Wetzel

## Abstract

1. For 10,000 years humans have altered plant traits through domestication and ongoing crop improvement, shaping plant form and function in agroecosystems. To date, studies have focused on how these processes have shaped whole-plant or average traits; however, plants also have characteristic levels of trait variability among their repeated parts, which can be heritable and mediate critical ecological interactions. As concerns about sustainable pest management increase, there is growing interest in approaches that increase trait diversity in crop agroecosystems. Here, we examine an under-appreciated scale of trait variation – among leaves, within plants – that may have changed through the process of domestication and improvement in a key crop.
2. We explore how levels of within-plant, among-leaf trait variability differ between cultivars and wild relatives of alfalfa (*Medicago sativa*), a key forage crop with an 8,000 year cultivation history. We grew individual plants from 30 wild populations and 30 cultivars, encompassing a range of domestication and improvement histories. For each plant, we quantify variability in a broad suite of physical, nutritive, and chemical leaf traits, including measures of chemical dissimilarity (beta diversity) among leaves.
3. We find that intra-individual trait variability has changed over the course of domestication and crop improvement, with effects often larger than changes in trait means. Cultivated alfalfa had elevated variability in SLA, trichomes, and C:N; increased diversity in defensive compounds; and reduced variability in phytochemical composition. We also elucidate fundamental associations between trait means and overall investment in secondary metabolites with patterns of among-leaf variability and chemical diversity.
4. We conclude that within-plant variability is an overlooked dimension of trait diversity in this globally critical agricultural crop. We find that trait variability is actually higher in cultivated plants compared to wild progenitors for multiple nutritive, physical, and chemical traits, highlighting a scale of variation that may mitigate loss of trait diversity at other scales in alfalfa agroecosystems and in other crops with similar domestication and improvement histories.

## Introduction

Humans exert powerful evolutionary forces on the plants we grow. Compared to wild relatives, modern plant cultivars often grow faster or larger, follow altered life histories, and differ in physical, nutritive, and chemical traits (Meyer *et al.*, 2012; Whitehead *et al.*, 2017). Understanding these trait differences is vital to human health and agroecological management (Wood *et al.*, 2015). To date, most studies of plant traits in domesticated cultivars have focused on trait means, such as mean nutritive or defensive content of fruits or leaves. However, this obscures a fundamental dimension of the plant phenotype: the level of trait *variability* among plant repeated structures. While this intra-individual variability is often dismissed as noise, studies in wild plants suggest it may be a key scale of diversity: it is often greater than variability among individuals and influences critical biotic interactions including herbivory and pollination (Herrera, 2009; Wetzel *et al.*, 2016; Pearse *et al.*, 2018). Moreover, recent work finds that within-plant variability can be heritable and controlled by loci independent of those shaping trait means (Bruijning *et al.*, 2020), or generated by processes common during domestication and improvement – such as inbreeding, polyploidization, or hybridization – suggesting that variability *per se* may be a common, yet overlooked aspect of modern plant cultivars. Indeed, differences in trait variability between crops and wild plants could represent yet another scale of agroecological biodiversity loss – or constitute an underappreciated scale of trait diversity that could be used to enhance ecosystem services (Wood *et al.*, 2015).

Domestic plants could have greater or lesser levels of trait variability compared to wild progenitors due an array of mechanisms, and as either a by-product of selection on other plant traits or as a direct target of crop improvement. Fixed relationships between trait means and variances, often positive (Herrera, 2009; Nakagawa & Schielzeth, 2012), could cause selection on traits means to indirectly change variability. This phenomenon could make selection for higher average nutrient content lead to an increase in nutrient variability among leaves, a surprising side effect of crop improvement. Similarly, relationships between the concentration and richness of phytochemicals (Wetzel & Whitehead, 2019) suggest that selection on overall phytochemical production could indirectly change phytochemical diversity within and among leaves. Selection on whole-plant traits, such as growth, could indirectly alter the range of traits achieved across tissue ontogeny (Fiorani *et al.*, 2000) or increase developmental instability and trait ‘noise’ as leaves mature (Arendt, 1997). Such developmental instability, and subsequent effects on levels of trait variation among plant tissues, could also increase with hybridization (Møller & Shykoff, 1999; Veličković & Stanković, 2005; Albarrán-Lara *et al.*, 2010) – a common part of the domestication and improvement process (but see Gardner, 1995; Waldmann, 1999). In contrast, genome duplication, another common feature of domesticated plants (Ramanna & Jacobsen, 2003; Salman-Minkov *et al.*, 2016), has been found to buffer and minimize stochastic gene expression (Cook *et al.*, 1998; Klingenberg, 2003; Soltani *et al.*, 2016). Alternatively, variability could be under direct selection; humans may have favored homogeneity within plants, which could facilitate harvesting and marketability (Liu *et al.*, 2005), or favored cultivars with greater variability – regardless of the underlying mechanism – if it deterred agricultural pests (Wetzel *et al.*, 2016; Pearse *et al.*, 2018), promoted less costly patterns of herbivory (Mauricio *et al.*, 1993), or allowed plants to cope with novel agricultural environments via enhanced phenotypic plasticity (Grossman & Rice, 2012) or adaptive noise (Viney & Reece, 2013) among tissues.

Despite the many mechanisms that could change within-plant trait variability over the course of domestication, we lack a basic understanding of the presence, strength, or direction of changes in variability, as well as their relation to trait means and tissue ontogeny. Such an understanding could advance agricultural sustainability by informing new ways to use trait diversity for management – a key focus of agroecology (Wood *et al.*, 2015) – at a scale of critical but often overlooked importance to ecological interactions (Real, 1981; Suomela & Ayres, 1994). Indeed, variability *per se* can reduce herbivore performance for traits that are both costly or beneficial in terms of their mean (Wetzel *et al.*, 2016; Pearse *et al.*, 2018) and, when large, can even supersede the associated mean as a driver of ecological interactions (Caraco & Lima, 1985). To date, fine-scale trait variability has been shown to influence performance of herbivores (Wetzel *et al.*, 2016; Pearse *et al.*, 2018) and levels of costly damage (Suomela & Ayres, 1994), inhibit pathogen spread (Zhu *et al.*, 2000) and alter pollination (Real, 1981; Herrera, 2009). Therefore, the goal of this work is to explore whether within-plant trait variability differs between domestic plants and their wild progenitors, and to compare the magnitude of change in variability to that of accompanying trait means. By documenting patterns of among-leaf variability across a broad suite of plant functional traits, our goal is to highlight variability *per se* as an important axis of the domesticated plant phenotype, and encourage future work into the mechanistic basis and ecological consequences of trait variation at this scale in this and other crop systems.

To achieve this goal, we compare levels of within-plant trait variability between modern cultivars and wild relatives of alfalfa (*Medicago sativa),* a key forage crop with an 8,000-year domestication history. As for most crop plants, identification of a single wild progenitor population or gene pool is challenging (Milla *et al.*, 2015) and, in the case of alfalfa, further complicated by multiple origins and complex hybridization and introgression events (Muller *et al.*, 2003). Therefore, we took the approach of comparing domestic plant genotypes with individuals from wild relative populations (Turcotte *et al.*, 2014; Whitehead *et al.*, 2017), selecting one genotype from each of 30 wild progenitor populations and each of 30 domesticated cultivars (Table S4). This allowed us to compare phenotypes from an extensive range of cultivar domestication histories and putative wild ancestor populations across the center of origin, rather than allocate replication within specific, and unknown evolutionary histories (Whitehead & Poveda, 2019). To measure intra-individual trait variability, we quantified the standard deviation of multiple nutritive, physical, and chemical traits and the beta diversity of phytochemical composition among 9 leaves per plant, stratified across leaf ontogeny (Fig. S1.1). Specifically, we ask (i) How does intra-individual, among-leaf trait variability change with domestication? (ii) Can changes in trait variability be explained by trait means, or are they an independent feature of the domesticated phenotype? (iii) To what degree do changes in trait variability reflect changes across leaf ontogenetic trajectories, versus pervasive shifts within age classes?

## Materials and Methods

### Selection of wild plant populations and cultivars

Modern alfalfa cultivars (*Medicago sativa*) trace their 8,000 year domestication history to three wild diploid subspecies (*M. sativa* ssp. *falcata, M. sativa* ssp. *caerulea* and their naturally-occurring hybrid, *M. sativa* ssp. *hemicycla),* with a potential role for several tetraploid subspecies (Havananda *et al.*, 2010). Like many crop species, modern alfalfa is the product of inbreeding (Kimbeng & Bingham, 1999; Katepa-Mupondwa *et al.*, 2002) [though moderate; see (Holland & Bingham, 1994)), extensive hybridization among subspecies and cultivars (Barnes, 1977; Maureira *et al.*, 2004), and also polyploidization (Capomaccio *et al.*, 2010) – all of which could contribute to levels of trait variability or homogeneity among tissues within plants (see above). We used this information to acquire a broad array of wild progenitor genotypes from across the native range of Vavilov’s “Near Eastern” and “Central Asiatic” centers of crop origin (Barnes, 1977), focusing on the three diploid subspecies (ssp. *falcata,* ssp. *caerulea,* and ssp. *hemicycla*) as the origin of tetraploid subspecies is less clear and may be linked to ongoing gene flow with cultivars, of which all are tetraploid (Fig S1.2; Table S1.4) (Small & Bauchan, 2011). We then selected 30 domestic cultivars encompassing a range of domestication histories, using information about original agricultural introductions of the plant (Barnes, 1977) (Fig S1.2; Table S1.4). The goal of this sampling was to represent a broad array of phenotypes among wild progenitors as well as among modern cultivars, given the long history of alfalfa cultivation, hybridization, and development and thus incorporating the many aspects of domestication and improvement that could shape levels of within-plant, among-leaf trait variability. See supplemental methods for further detail on plant selection.

### Common garden

In late May 2018 we imbibed and germinated 15-20 alfalfa seeds/population or cultivar on filter paper, following scarification (100-grit sandpaper) and a bleach soak (3% solution for 10min). Seedlings were established in cell trays of standard peat mix in the greenhouse. After 15 days, we transplanted N = 240 healthy plants (N = 4/population) into individual pots containing a mix of peat mix and field soil (50%-50%), to acclimate them to field soil conditions. In early July, after ca. 30 days of growth in the greenhouse, we outplanted one individual per population/cultivar (N = 60) into a common garden at Kellogg Biological Station (Hickory Corners, MI). Plants were arranged in a stratified design, alternating positions between wild and domestic populations. Within this ‘checkerboard’ layout, populations or cultivars were randomly assigned to each position. We placed a parallel set of plants (N = 60) into screenhouses next to the common garden, to experience similar abiotic conditions as field plants, but protected from herbivores to avoid differential induction of secondary chemistry between wild and domestic plants.

### Trait measurement

We collected leaf tissue for physical (specific leaf area, trichome density) and nutritive (leaf water content, carbon and nitrogen content) traits in August of 2018 from plants in the common garden (64 days of growth). To do so, we randomly selected 3 vegetative stems (peduncles) per plant individual. From each stem, we selected the expanding leaf at the tip (“young”), the expanded leaf at the middle nodal position of the peduncle (“middle”), and the basal leaf, where the peduncle joins the main stem (“old”) for a total of 9 individual leaves per plant (See Fig. S1.1). We avoided leaves with evidence of physical damage, pathogens or herbivory. Following removal, each leaf was immediately weighed, scanned, photographed under magnification, and dried in a coin envelope. We quantified SLA, LWC, and trichome density for each leaf; elemental analysis of nitrogen and carbon was assessed by flash combustion by the University of Georgia Stable Isotope Ecology Laboratory (see supplemental methods for details of trait measurement). At the time of leaf collection, we also measured plant height and width for a measure of plant size (volume).

Due to logistical constraints in 2018, we quantified variability in chemical traits in the following year, using a second set of plants (see above). We followed the same protocol described above to harvest 9 leaves per plant from individuals in screen cages (see above). All samples were dried in a drying oven; weighed, homogenized using a bead mill (60sec), and extracted individually using EtOH solvent containing a 100 nm digitoxin as the internal standard. Because leaf masses varied by orders of magnitude, we adjusted the solvent volume in proportion to the tissue mass (1mg : 300uL solvent), rather than choosing a single (by necessity, very low) mass to add across all vials. As an additional “overall” reference, a pooled sample was prepared by adding an equal volume of all extracts to one vial, to be used for quality control (monitoring retention time stability and signal intensity during runs) and for compound identification purposes. We also prepared solvent blanks containing 100nM EtOH and digitoxin solution. Ethanolic leaf extracts were analyzed by LC/MS using a Thermo Q-Exactive mass spectrometer interfaced with a Thermo Vanquish UPLC system. Data were acquired using a FullMS method for quantitative analysis and a data-dependent MS/MS (DDA) method for generating MS/MS spectra for compound identification. To reduce prevalence of false positives in the data, we converted all raw peak areas below 10000ppm to zero, as this is near the instrumental detection limit. We then normalized compound peak areas to the peak intensities of digitoxin in each sample. Lastly, to reduce chemical noise in the data, we subtracted the mean peak area found in blanks (N = 29 blanks) from their peak areas quantified in each sample. We focused on identification of triterpene glycoside saponins compounds, a major class of compounds in alfalfa of known resistance function against herbivores (Nozzolillo *et al.*, 1997). See supplement for further description of plant chemistry methods.

### Quantifying variability and statistical analysis

For each trait, we used three model structures to explore how variability shifts with domestication, and the degree to which shifts occur jointly with or independently from changes in trait means. Specifically, we asked 1) How does trait variability shift with domestication (*σ_χ_* ~ *D*); 2) How do trait means shift with domestication 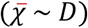; and 3) Do shifts in variability occur independently of, or jointly with, shifts in trait magnitude 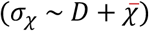. By comparing domestication effects across these three models, we can quantify overall shifts in both attributes of the plant phenotype, as well as the degree to which these trait outcomes occur independently or jointly.

For physical traits (SLA, trichome density), nutritive traits (LWC, C and N content, C:N ratio), and individual saponin compounds we used the standard deviation and mean trait values among leaves. For chemical diversity, we quantified richness, Shannon’s, and Simpson’s diversity of all saponins (N = 86) (function *diversity,* R package “vegan”, version 2.5-6) (Oksanen *et al.*, 2019) within individual leaves to represent alpha chemical diversity (α); pooled across all leaves as a measure of gamma diversity (γ); and Bray-Curtis dissimilarity in presence/absence, proportional abundance, and raw abundance among leaves (functions *vegdist* and *betadisper,* R package “vegan”, version 2.5-6) (Oksanen *et al.*, 2019) to quantify beta diversity (β). For *α, γ,* and β chemical diversity, we used the peak area at the corresponding scale to represent total compound production, and parallel use of the mean for other traits (summed concentration per leaf for α diversity; average across all leaves for *γ* diversity, and average per-leaf concentration among all leaves or within leaf age class for β diversity at the whole-plant and within age-class scales, respectively). In this way, we can ask how multivariate aspects of chemical diversity are related to the overall quantity of compounds produced by plant leaves. All metrics of physical, nutritive, and chemical variability and means were calculated at the whole-plant scale (across N = 9 leaves of all ages) as well as within young, middle, or older-aged leaves per plant (N = 3 leaves/stage). Together, these ten diversity measures describe multiple dimensions of within-plant trait variability, generated by differences in trait amount as well as in trait identity among and within leaves of differing ontogenies.

We used generalized linear mixed models (GLMMS) to test for differences in trait means and within-plant variability between wild and domestic plants. All models were implemented using the function *glmmTMB* (R package “glmmTMB”, version 1.0.0) (Brooks *et al.*, 2017) in the statistical computing environment *R* (version 3.6.0) (R Core Team, 2019). For all models, we used domestication status (wild, domestic) as a main effect to predict each response variable at two scales: among all leaves (N = 9 leaves/plant) and within each leaf age class (N = 3 leaves/plant/class). For analyses within leaf age class, we used age (young, middle, old) as an additional main effect, and tested for interactions with domestication status. We also used plant individual as a random effect in these models, as each plant yielded three trait measures (one per age class). For analyses of individual compound variability (SD) among leaves, we included compound identity as a random effect, and allowed the relationship between mean and SD to vary within each compound as well within each plant individual (random slopes). For analyses of variability across ontogeny, we excluded N = 2 compounds that were not found across all leaf stages, as variability values would fail to represent comparable ranges of leaf ontogeny. For analyses of compound diversity (richness, Shannon), we allowed the relationship between diversity and total peak area to vary within plant individual (random slopes). For each analysis, we performed model selection using AIC (function *dredge*; R package “muMIn”, version 1.43.6 and function *AICctab*; R package “bbmle”, version 1.0.20) (Bartoń, 2016; Bolker & R Development Team, 2017).

To account for differences in plant growth rate, and the potential role of this trait in shaping levels of among-leaf trait variability, we used plant size (Fig. S1.5) as a covariate in all models. In the absence of genetic information for all cultivars and wild populations, we also included the estimated contribution of wild subspecies (Barnes, 1977) to each plant population or cultivar (Table S1.4) as potential covariates during model selection. In this way, we incorporated the non-independence of cultivars with greater or lesser contributions of each wild subspecies in their breeding history. See Tables S2.1 and S2.2 for full model structures, and terms included in top models for each trait.

To compare differences between wild and domesticated plants across traits with such a diverse array of measurement units (e.g. from trichome densities to Bray-Curtis distances), we converted all effect sizes to percent change in response variables (means or variability) with domestication. For each trait and model structure, we obtained 95% CIs for the effect of domestication using the function *bootMer* (R package “lme4”; nsim=500) (Bates *et al.*, 2015). We then used the *predict* function (R Core Team, 2019) implemented via the *ggpredict* function (R package “ggeffects”, version 0.13.0) (Lüdecke, 2018) to calculated the corresponding percent changes associated with beta and its 95% CI. Thus, positive % change values represent an increase with domestication, and negative values represent a decline with domestication. We obtained p-values for domestication effects using the *Anova* function (R package “car”, version 3.0-6) (Fox *et al.*, 2011). For post-hoc contrasts (e.g. effects of domestication within leaf age class), we conducted t-tests for using the function *pairs* (R package “emmeans”, version 1.4.1) (Lenth, 2019). Data visualization was performed using ggplot2 (version 3.2.0) (Wickham, 2016).

## Results

We found that domestic cultivars exhibited greater intra-individual variability in physical and chemical traits in alfalfa (Fig. 1b; Table S1.1). Effect sizes were large; relative to wild plants, the trait variability encompassed by domestic plant leaves across ontogeny was 83.9% greater for specific leaf area (SLA) (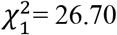, p<.001), 37.3% greater for the concentration of individual saponins (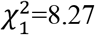, *p*<.01), and 43.3% greater for the summed concentration of all saponins (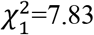, *p*<.01) (Fig1b). In contrast, we observed no differences in among-leaf variability in trichome density, leaf water content, carbon, nitrogen, or C:N ratio between domestic and wild plants at this scale. Chemical diversity also increased with domestication: the richness of saponins encompassed by domestic plant leaves (*γ*) was 6.6% greater (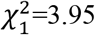, *p*<.05) than in wild plants. In contrast to *γ* diversity, variability in multivariate chemical composition among those leaves (*β* diversity) was 13.7% lower in domestic plants (proportional compound abundances: 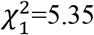, *p*=.02) (Fig. 1b).

**Figure 1.**
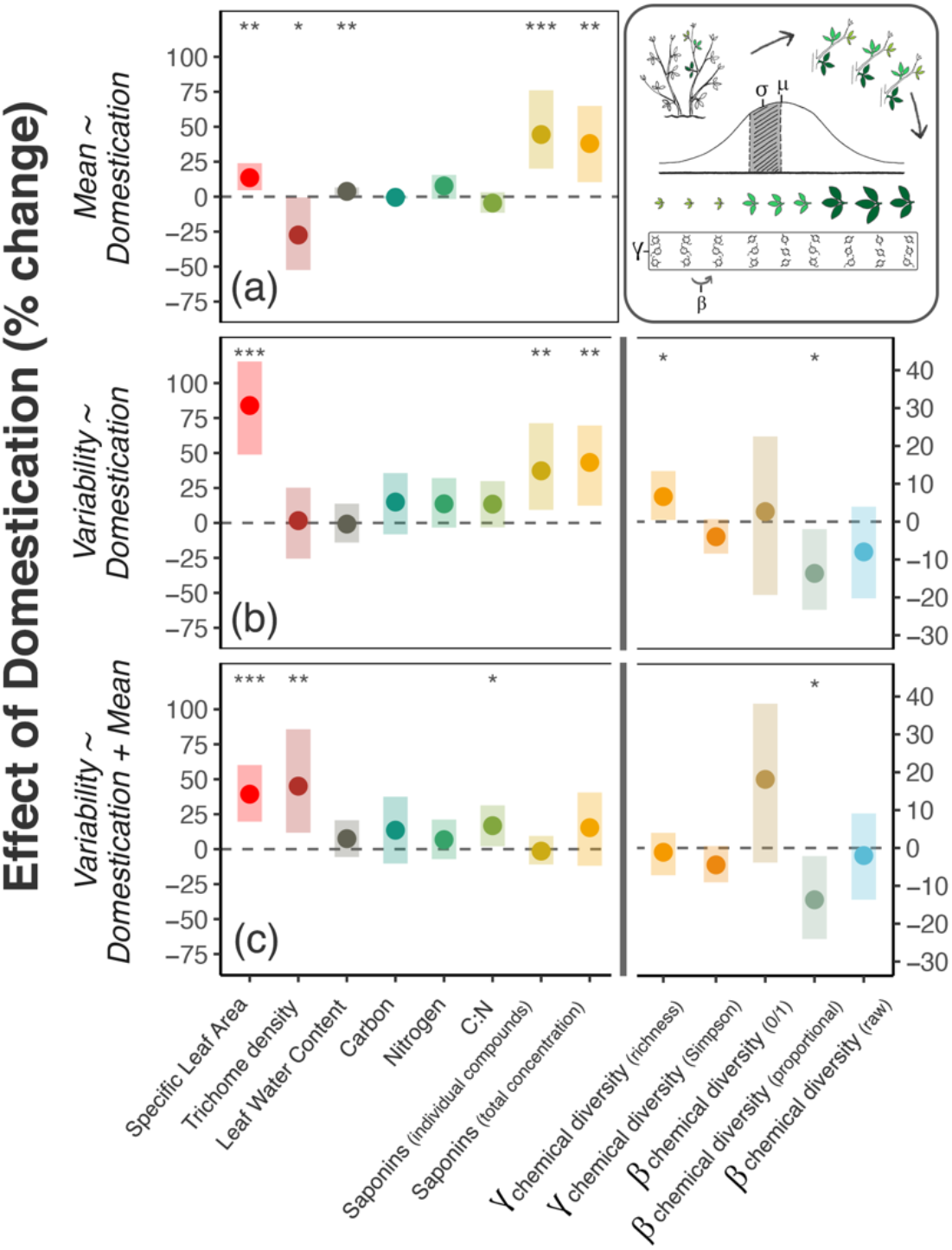
Comparing levels of among-leaf, within-plant trait variability between domestic cultivars and wild relatives of alfalfa, and relationship with trait means. Differences in within-plant trait variability and trait means between wild and domestic alfalfa plants. Points are model-estimated domestication effects (% change), and shaded bars are the 95% CIs. Top panel (a) shows change in trait means and total compound concentrations, averaged across 9 leaves per plant; middle panel (b) shows change in trait variability (standard deviation) and chemical diversity across the same 9 leaves per plant; and bottom panel (c) shows the change in the same among-leaf variability traits, but after accounting for mean ~ variance and compound concentration ~ diversity relationships. Note two different y-axis ranges; righthand y-axis is for chemical diversity traits. Asterisks show model significances at the <.05, <.01, and <.001 levels. Top right: schematic of sampling design within each plant and metrics of variability (see also Fig. S1).

Next, we asked whether differences in trait variability between wild and domestic plants could be explained by changes in trait means or total compound concentrations. Consistent with the literature on plant domestication (Whitehead *et al.*, 2017), we observed change in the means and total levels of multiple physical, nutritive, and chemical traits (Fig. 1a; Table S1.2). Shifts in trait averages among leaves explained changes in variability for some traits, but for others variability changed independently of or opposite to changes in means. For example, greater variability in individual saponin compounds was explained by their 44.4% higher concentrations, on average, in domestic plants (Fig. 1 a & c; Table S1.2-S1.3), and greater total compound production (+38.0%) explained the higher *γ* richness among domestic plant leaves (Fig. 1c; Table S1.2-3). In contrast, SLA variability was still 39.3% greater among leaves of domestic plants (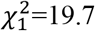, *p*<.001), even after accounting for shifts in average SLA with domestication (Table S1.3); trichome variability was 45% greater (t_55_=−2.81, *p*<.01) after accounting for a 27.4% decline in mean densities (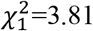, *p*=.05); and among-leaf variability in C:N ratios increased by 16.8% with domestication (t_38_=−2.21, *p*<.05), while average C:N ratio did not differ (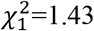, *p*>.1) (Fig 1c; Table S1.3). Among-leaf β diversity in proportional saponin concentrations was still 13.7% lower among domestic plant leaves, after taking overall compound production into account (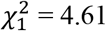, *p* < .05) (Fig 1c; Table S1.3).

To understand how differences in trait variability between wild and domestic plants are related to leaf ontogeny, we quantified trait mean and variability among leaves in early, middle, and later stages of expansion (Fig. 2a & b). We found that effects of domestication on variability were also observed within leaf age classes, as well as across leaf ontogeny. For example, variability in SLA and LWC increased 18.8% and 17.2 %, respectively, among leaves within each age class (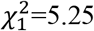, *p*<.05 and 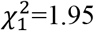, *p*<.01), after accounting for changes in means between wild plants and cultivars (Fig. 2a & c, Table S1.3). Differences in saponin concentration variability between wild and domestic plants depended on leaf age class, reaching a 93.3% increase in total concentration variability for young leaves of domestic compared to wild plants (*t*_170_=−3.10, *p*<.01; Table S1) – larger than the 76.0% increase in the corresponding mean (*t*_170_=−3.56,p<.001; Table S1.2). Chemical richness and Simpson’s diversity were 8.5% higher and 3.0% lower, respectively, for young, middle, and older leaves (Fig 2b; Table S1.1), with effects associated with greater compound production in parallel to results among all leaves (Fig 2c; Table S13). β chemical diversity in proportional concentrations declined by 16.1% / 19.4% (with / without total compound production as a covariate) from wild to domestic plants in each leaf age class (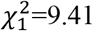9.41, *p*<.01; 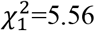, *p*<.05, respectively) – an effect size of greater magnitude than that quantified across leaf ontogeny (Table S1.3). For other traits, differences in variability between wild and domestic plants occurred within one age class but not others (C:N ratio; Fig. 2b & c, Table S1.3).

**Figure 2.**
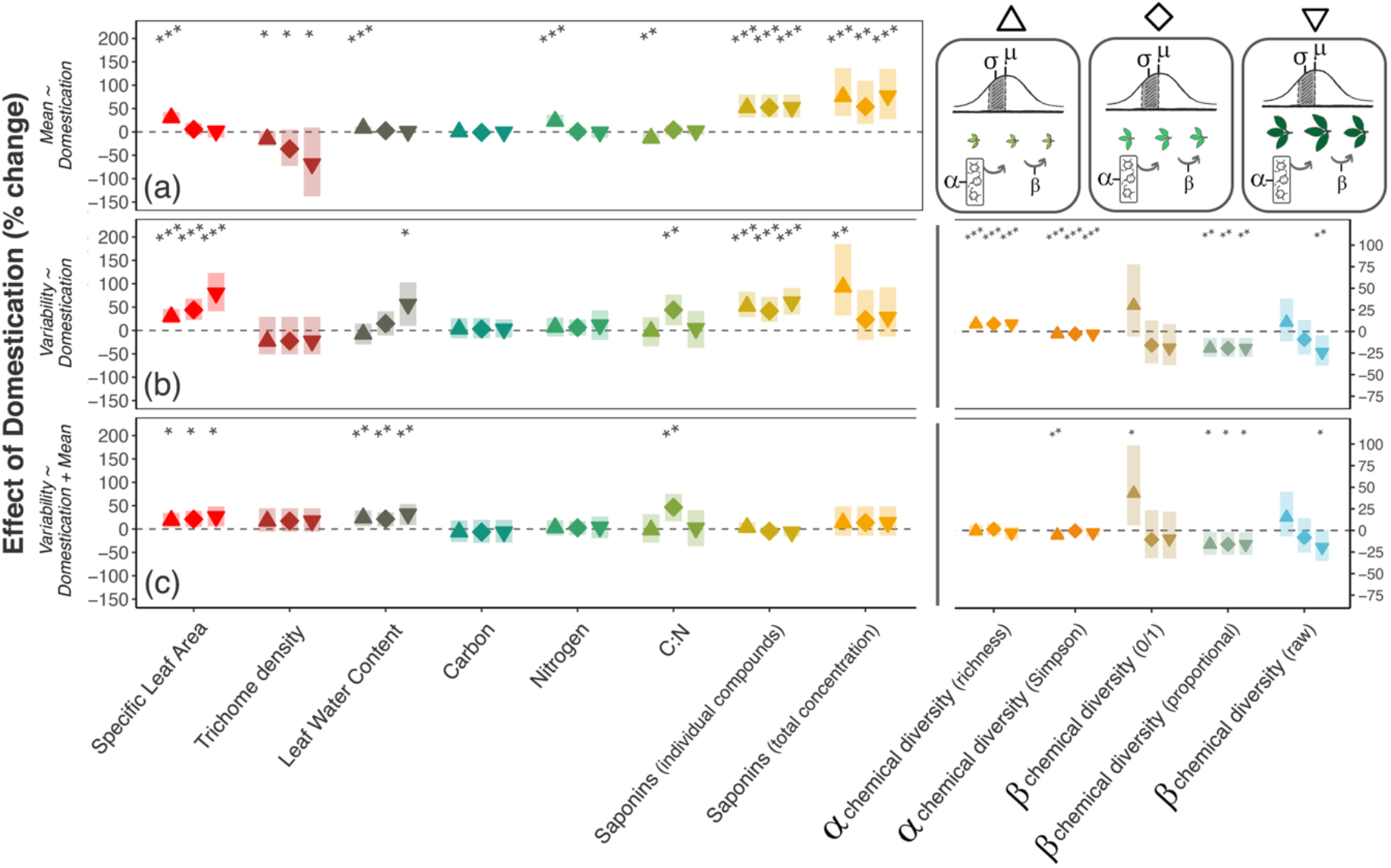
Differences in trait variability between wild and domestic plants depend on leaf ontogeny. Certain leaf age classes show more pronounced effects of domestication on among-leaf trait means or variability. Points are model-estimated domestication effects (% change), and shaded bars are the 95% CIs. Point shape indicates leaf age class: young/expanding 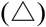, expanded 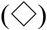, and older/expanded 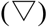. Top panel (a) shows change in trait means and total compound concentrations, averaged across 3 leaves per plant, per age class (young, middle, old); middle panel (b) shows change in trait variability (standard deviation) and chemical diversity across the same 3 leaves per plant/age class; and bottom panel (c) shows the change in the same among-leaf variability traits, but after accounting for mean ~ variance and compound concentration ~ diversity relationships at this age-specific scale. Note two different y-axis ranges; righthand y-axis is for chemical diversity traits. Asterisks show model significances at the <.05, <.01, and <.001 levels. Top right: schematic of sampling design within each plant and metrics of variability (see also Fig. S1).

Wild and domestic plants differed in plant size – a proxy for growth rate – with domestic plants attaining 34.5cm^2^ larger volume than their wild relatives (Figure S1.5). Plant size was not a strong predictor of traits; it was rarely included in top models during model selection and, when it was included, effect sizes were small and generally not significant (Tables S2.1, S2.2). The contribution of wild subspecies was often selected in top models (either ssp. *caerulea* or ssp. *falcata*), and was sometimes significantly associated with trait mean values or levels of variability: for example, plants with greater percentage ssp. *caerulea* tended to have lower Simpson’s diversity, and those with more ssp. *falcata* tended to have greater average Nitrogen and lower C:N ratios among all leaves; however, overall effect sizes were small and, similarly to plant size, were often not significant (Tables S2.1, S2.2).

## Discussion

We find that wild and domestic plants differ not only in the average value of plant functional traits among leaves, but also the magnitude of among-leaf variability within plant individuals. Specifically, we find that domesticated *M. sativa* have increased variability in leaf physical and nutritive traits and concentrations of individual saponin compounds, and a restructuring of chemical diversity from greater among-leaf diversity in wild plants toward greater within-leaf diversity (e.g. alpha diversity) in domestic plants – independent of shifts in trait means or overall compound production. As the importance of trait variability *per se* to a range of ecological process is increasingly acknowledged (Herrera, 2009; Wetzel *et al.*, 2016; Pearse *et al.*, 2018), our study highlights this component as an underappreciated facet of the domesticated *M. sativa* phenotype and, potentially, an overlooked source of trait variability across a range of crop systems.

Consistent with previous studies of plant trait evolution with domestication, we find frequent differences in the averages of many leaf nutritive, physical and chemical traits of domestic plants and their wild progenitors (Delgado-Baquerizo *et al.*, 2016). Importantly, we find that these domestication effects differ in magnitude and prevalence with leaf age, suggesting that selection may be shaping the ontogeny of leaf functional traits. We find changes in average physical and nutritive traits known to mediate photosynthetic capacity and herbivore damage: for example, domestic *M. sativa* have higher SLA than wild relatives, perhaps indicating selection for greater photosynthetic area at the cost of tissue robustness and resistance to herbivores (Westoby *et al.*, 2000). Average trichome densities declined sharply with domestication, while leaf water content and nitrogen content increased, particularly among leaves earlier in expansion. We also observed that domestic plants produce much higher concentrations of secondary metabolites than their wild progenitors, both for individual saponin compounds as well as their total concentrations. Together, these shifts in overall trait production and investment are consistent with selection for growth-related processes (Fig S1.5) and forage nutritional quality (Delgado-Baquerizo *et al.*, 2016) shaping vigorous domesticated phenotypes that excel in both primary and secondary metabolism. Such positive associations between plant growth rate and defensive production are often observed among plant individuals, particularly for species that have evolved in more resource-rich environments (Hahn & Maron, 2016), and is consistent with meta-analyses finding both increases and declines in secondary chemistry and other defense traits with domestication (Meyer *et al.*, 2012; Turcotte *et al.*, 2014; Whitehead *et al.*, 2017).

In addition to shaping different overall investment in physical, nutritive, and chemical traits, we find that levels of trait variability among *M. sativa* plant leaves have also changed over the course of domestication and improvement– with differences often of greater effect size than those of trait averages. Variability in trichome densities, specific leaf area, leaf water and C:N content all increased from wild to domestic plants – even as their averages declined, increased, or showed no effect: thus, modern *M. sativa* cultivars show a general increase in variability of physical and nutritive traits, independent of strength or direction of shifts in the mean as compared to wild relatives. In contrast, greater chemical production over the course of domestication shapes variability of individual saponins among leaves and leaf-level compound richness, but does not explain the effects of domestication on variability in compound composition. Thus, while the individual saponins in domestic plants differ more in their concentrations among leaves, they retain more consistent abundances relative to each other; in contrast, saponins in wild plants vary less in their absolute concentration but more in their ratios, increasing among-leaf variability in chemical composition. As compound ratios can shape synergistic vs. antagonistic outcomes (Caesar, 2019), this result suggests that wild plants maintain potential for a greater variety of biological effects on consumers among their leaves, while domestic plants confront herbivores with a similar ‘cocktail’ from leaf to leaf. The multi-scalar prevalence of these differences between wild and domestic plants – from across leaf ontogeny to equally or more pronounced effects within certain leaf age classes - suggest that shifts in variability are not simply due to greater trait maxima or lower trait minima reached across leaf ontogeny in cultivars; rather, altered trait variability is a pervasive feature of domestic *M. sativa* leaves, with more pronounced differences between wild and domestic plants at some stages of leaf development than others. Thus, herbivores that specialize within leaf age class may experience different domestication effects, and tissue-age generalists may find leaf age or nodal position to be a noisier quality cue.

More broadly, the complex relationships between trait magnitude and variability observed in this system suggest that crop domestication can shape within-plant variability via multiple pathways and processes. If anthropogenic selection has directly targeted trait averages, or shaped allocation via tradeoffs between growth and defense, we should look for corresponding shifts in levels of trait variability and chemical diversity. However, for some traits these two properties may be de-coupled, suggesting that within-plant variability could be manipulated independently of trait means. These results parallel recent studies finding both independent and shared genetic control of trait means and their variability within individuals (Bruijning *et al.*, 2020). Our results are also consistent with compromised leaf development through the process of domestication and improvement. Previous studies find that processes involved in domestication and improvement – such as polyploidization – can increase trait ranges at other scales, such as among plant individuals (Baker *et al.*, 2017), and either exacerbate (Comai *et al.*, 2000; Madlung *et al.*, 2012) or mediate (Cook *et al.*, 1998; Klingenberg, 2003; Soltani *et al.*, 2016) developmental instability at the within-individual scale. Our findings are consistent with genome duplication in *M. sativa* resulting in poor developmental control as leaf traits progress through ontogeny, as well as other processes working to increase variation in trait expression among leaves – such as loss or disruption of coadapted gene complexes during hybridization (Graham & Felley, 1985; Siikamäki, 1999). As polyploidization and hybridization are ubiquitous features of the domestication and improvement process in many cultivated plants, we suggest that these findings may be mirrored across many crops beyond alfalfa.

Considering trait mean and variability among leaves as interactive attributes of the plant phenotype is of critical ecological relevance; variability *per se* can act to reduce herbivore performance for traits that are either harmful or beneficial in terms of their average (Wetzel *et al.*, 2016; Pearse *et al.*, 2018), and the importance of trait means or variability to consumers may depend on the magnitude of the other (Caraco & Lima, 1985). Thus, understanding among-leaf distributions in traits and chemical similarity, in addition to their means and overall compound production has critical implications for plant breeders balancing crop improvement with vulnerability to pest attack. For example, among-leaf variability or ‘noise’ in functional traits could provide a bet-hedging strategy, countering unpredictable abiotic or biotic conditions across leaves (Viney & Reece, 2013). For small consumers that move among leaves, encountering greater trait variability could reduce herbivory by introducing greater uncertainty during foraging (Real, 1981), requiring frequent physiological adjustments, and establishing costly chemical synergies (Wetzel & Thaler, 2016) – or increase herbivory by allowing performance-enhancing dietary mixing (Moreau *et al.*, 2003) or promoting chemical antagonisms that inhibit compound defensive function (Feng & Shoichet, 2006; Caesar, 2019). As many insect herbivores complete development within one or few plant individuals, trait variability at the within-plant scale may be particularly critical in shaping spatial patterns of damage within plants and associated costs to plant growth (Mauricio *et al.*, 1993), as well as frequency of risky herbivore movement (Real, 1981).

Our findings suggest that, over thousands of years of domestication and improvement, humans have not only been altering leaves on average – but also shaping the magnitude of trait variability around these means in a key crop plant. This finding, plus our growing appreciation for the ecological and evolutionary importance of variability *per se,* indicates that variability may be a key component missing from our understanding of domestication and agroecology more broadly. Indeed, modern agroecosystems have artificially low trait diversity at the landscape, community, and population scale, with cascading effects on plant damage, yield, and agroecosystem function (Wood *et al.*, 2015). Our work reveals a previously overlooked scale at which trait diversity differs between agricultural and natural systems: among leaves within plants. At the same time, the result that domestic cultivars may host greater variability for some traits, and in various association with trait means, suggests that the within-plant scale may be a novel frontier of trait diversity that could, through cultivar selection or plant breeding, be harnessed to mitigate low diversity at other scales and enhance sustainable pest management.

## Acknowledgements

We thank Sydney Jackson and Kalin Bayes for data collection; Mark Hammond, Brook Wilke, Brian Irish, and Zsofia Szendrei for feedback and logistics; Anurag Agrawal, Matthew Forister, Jeffrey Ross-Ibarra, Sharon Strauss, Gregg Howe, and Martin Turcotte for comments; Casey Philbin for chemistry discussions; and the Wetzel Lab for feedback and support. This work funded by USDA NIFA Postdoctoral Fellowship #12432349.

## Author Contributions

MR: conceptualization; methodology; analysis; writing; and funding acquisition. WW: conceptualization; methodology; editing; supervision. AS: methodology; analyses; editing. All authors contributed to the development of the manuscript, and declare no competing interests.

## Data Availability

Data and code will be made available at www.figshare.com. Raw LC-MS data will be made available through MetaboLights.

