## Appendix S1 for "A domestic plant differs from its wild relative along multiple axes of within-plant trait variability and diversity"

**Supplementary Information for**

**Title:** A domestic plant differs from its wild relative along multiple axes of within-plant trait variability and diversity

**Authors:** Moria L. Robinson[1,2,5]*, Anthony L. Schilmiller[3], William C. Wetzel[1,2,4,5,6]

[1]Department of Entomology, Michigan State University

[2]Kellogg Biological Station, Michigan State University

[3]Mass Spectrometry and Metabolomics Core, Michigan State University

[4]Department of Integrative Biology, Michigan State University

[5]Ecology, Evolution, and Behavior Program, Michigan State University

[6]AgBioResearch, Michigan State University

**This supplement includes:**

Supplementary text
Figures S1.1 to S1.5
Tables S1.1 to S1.5
SI References

**Supplementary Text**

*Selection of wild plant populations and cultivars.* Alfalfa is one of the most important agricultural plants in the world, grown across the globe for both animal forage and human consumption. It also boasts one of the longest and most complex domestication histories of any forage crop, having been cultivated by humans for over 8,000 years (Russelle, 2001) from a species complex with few barriers to gene flow (Bagavathiannan & Acker, 2009; Şakiroğlu & Brummer, 2011). Studies of alfalfa cultivars trace their domestication history to three wild subspecies (*M. sativa* ssp. *falcata*, *M. sativa* ssp. *caerulea* and their naturally-occurring hybrid, *M. sativa* ssp. *hemicycla*; diploid), with a potential role for several tetraploid subspecies (e.g. *M. sativa* ssp. *varia; M. sativa* ssp. *sativa*) that may have arisen in the wild or through hybridization with cultivars (Havananda *et al.*, 2010). We used this information to acquire as broad an array of wild progenitor genotypes as possible, from across the native range of Vavilov's "Near Eastern" and "Central Asiatic" centers of crop origin (Barnes, 1977). We focused on the three diploid subspecies (ssp. *falcata*, ssp. *caerulea*, and ssp. *hemicycla*), as the origin of tetraploid subspecies is less clear and may be linked to ongoing gene flow with cultivars or, if derived from evolution in nature, are also hypothesized to have arisen from among those diploid progenitors (Small *et al.*, 1990; Bagavathiannan & Acker, 2009; Havananda *et al.*, 2010) (Fig S1; Table S4). We then selected 30 domestic cultivars encompassing a range of domestication histories, both pre-and post-arrival to the United States, using information about original agricultural introductions of the plant (Barnes, 1977) (Fig S1; Table S4). The goal of this broad sampling was to represent a comprehensive array of phenotypes among wild progenitors as well as among modern cultivars, given the long history of alfalfa domestication, hybridization, and cultivar development, and incorporate the many aspects of domestication and improvement that could shape levels of within-plant, among-leaf trait variability.

*Trait measurement – nutritive and physical traits*

Each leaf was immediately weighed using an analytical balance (Mettler Toledo MS104TS), scanned, photographed under magnification, and dried in a coin envelope. We used ImageJ image analysis software to quantify leaf area and trichome density (Rasband, 1997). Specific Leaf Area (SLA) was calculated following the formula $SLA = \frac{leaf\ area_{(fresh)}}{leaf\ mass_{(dry)}}$. We quantified trichome density using a transect method, counting the number of intersections per mm on a line perpendicular to the midrib, using the middle leaflet of each trifoliate leaf. Leaf water content was quantified as $LWC = \frac{leaf\ mass_{(fresh)} - leaf\ mass_{(dry)}}{leaf\ mass_{(fresh)}} \times 100$. Elemental analysis of nitrogen and carbon content was assessed by flash combustion by the University of Georgia Stable Isotope Ecology Laboratory. To prepare samples for elemental analyses, we manually pulverized each leaf using a razor blade and placed 0.5-3.0mg of homogenized tissue into aluminum capsules. Manual pulverization, rather than use of a bead mill, was required due to the small size of leaves.

*Trait measurement – phytochemistry*

Leaves were removed directly from plants, using the method described above to stratify among age classes (N = 9 leaves/plant, across 3 relative age classes), and placed in individual coin envelopes. We also collected a second set of 9 leaves as a single aggregate/bulked sample per-plant, to serve as additional references for compound identification. We focused on identification of saponin compounds, due to their known resistance function against herbivores (Nozzolillo *et al.*, 1997; Wilding). All samples were then dried in a drying oven (no heat; air circulation only). In the lab, leaves were weighed, homogenized using a bead mill (60sec), and extracted individually using EtOH solvent containing a 100 nm digitoxin as the internal standard. For the set of bulked leaves, all leaves were homogenized together. Vials were agitated at room temperature (23C) for 10min to ensure contact between solvent and tissue in a ThermoMix (23C). Because leaf masses varied by orders of magnitude, we adjusted the solvent volume in proportion to the tissue mass (1mg : 300uL solvent), rather than choosing a single (by necessity, very low) mass to add across all vials. If leaves were > 7mg, they were homogenized with a razor blade and then subsampled to < 7mg, to remain within the capacity of the extraction vial. Following extraction, vials were centrifuged (10 min @ 13000 rcf) and 100 uL of supernatant was transferred to an LC-MS vial containing a small-volume insert. As an additional "overall" reference, a pooled sample was prepared by adding an equal volume of all extracts to one vial, to be used for quality control (monitoring retention time stability and signal intensity during runs) and for compound identification purposes. We also prepared solvent blanks containing 100nM EtOH and digitoxin solution.

*LCMS analysis.* Ethanolic leaf extracts were analyzed by LC/MS using a Thermo Q-Exactive mass spectrometer interfaced with a Thermo Vanquish UPLC system. 10 uL of sample was injected onto a Waters Acquity BEH-C18 UPLC column (2.1 x 100 mm) maintained at 40C. Compounds were separated using a gradient as follows: initial conditions were 95% solvent A (10 mM ammonium formate, pH 3.0; adjusted with formic acid) and 5% solvent B (acetonitrile containing 0.1% formic acid) and held until 0.5 min, ramp to 50% B at 2 min, ramp to 99% B at 7 min and then hold at 99% B until 8 min, return to 5% B at 8.01 min and hold at the initial conditions until 10 min. Ionization conditions were as follows: electrospray ionization in positive ion mode, capillary voltage was 3.5 kV, capillary temperature 256C, sheath gas 47.5, auxiliary gas 11.25, probe heater temperature was 412C, S-lens RF level was 50. Data were acquired using a FullMS method for quantitative analysis and a data-dependent MS/MS (DDA) method for generating MS/MS spectra for compound identification. The FullMS method used a scan range of m/z 120-1800, resolution setting of 70,000, AGC target of 3e6 and max ion time of 200 ms. The DDA method consisted of a survey scan from m/z 120-1800 at 35,000 resolution, AGC target 3e6 and max ion time of 100 ms; the top 5 ions were selected for MS/MS with an isolation width of 1.2 Da, resolution at 17,500, AGC target of 1e5, max ion time of 50 ms, and dynamic exclusion setting of 3 seconds. Solvent blank and pooled samples were run every 22 samples using the FullMS method. In addition the pooled sample and multi-leaf samples were run using the DDA method.

*Peak processing.* Non-targeted data analysis was performed using Progenesis QI software (Nonlinear Dynamics). Peaks were aligned across all samples but peak picking and deconvolution was done using only the data from ethanol blank samples, multi-leaf samples, and pooled sample to reduce processing time. An absolute ion intensity threshold of 10,000 and minimum chromatographic peak width of 0.1 minutes was set and peaks with retention time after 7.5 minutes were excluded from the analysis. Triterpene glycoside saponins are a major class of compounds in alfalfa and were identified based on accurate mass and MS/MS spectra present in the DDA data. Due to the large number of samples, the LC separation method was limited to 10 minutes to reduce instrument time and cost; however, this resulted in some poorly-resolved saponin peaks that Progenesis was unable to pick correctly. To generate accurate peak areas for these compounds, a second targeted analysis method was performed using Quanlynx in the Masslynx software (Waters).

*Normalization and filtering of peak areas.* To reduce prevalence of false positives in the data, we converted all raw peak areas below 10000ppm to zero, as this is near the instrumental detection limit. We then normalized compound peak areas to the peak intensities of digitoxin in each sample. Lastly, to reduce chemical noise in the data, we subtracted the mean peak area found in blanks (N = 29 blanks) from their peak areas quantified in each sample.

### Supplemental Figures

**Figure S1.1. Quantifying within-plant variability across multiple axes of the domestic plant phenotype.** We grew a single individual from each of N= 30 wild populations and N=30 cultivars of alfalfa (*Medicago sativa*). We emphasized breadth across domestication histories, rather than characterization of this trait for any particular wild population or cultivar, as our goal was to estimate levels of within-plant variability across as great a breadth of wild progenitor and domestic phenotypes as possible. This followed a design similar to other work (Whitehead & Poveda, 2019). Leaf sampling leveraged the modular structure of alfalfa plants; the multiple main stems produce lateral stems called 'peduncles', each of which is characterized by multiple leaves at various stages of expansion, from a single older basal leaf that defines the base of the peduncle to young expanding leaves at the apex. We used this organization to select leaves at differing stages of expansion: for each plant we selected three peduncles, and from each peduncle we collected the expanding leaf at the apical tip of the peduncle (early in expansion but with all three trifoliate leaflets visible); the leaf at the middle node, and the basal leaf (largest/fully expanded). We processed each leaf sample individually for physical, nutritive, and chemical traits. We quantified trait means, their associated variability (standard deviation), and measures of chemical diversity (alpha, beta, and gamma) at the whole-plant scale (N=9 leaves/plant) and within leaves in each age class (N = 3 leaves /age class/plant).

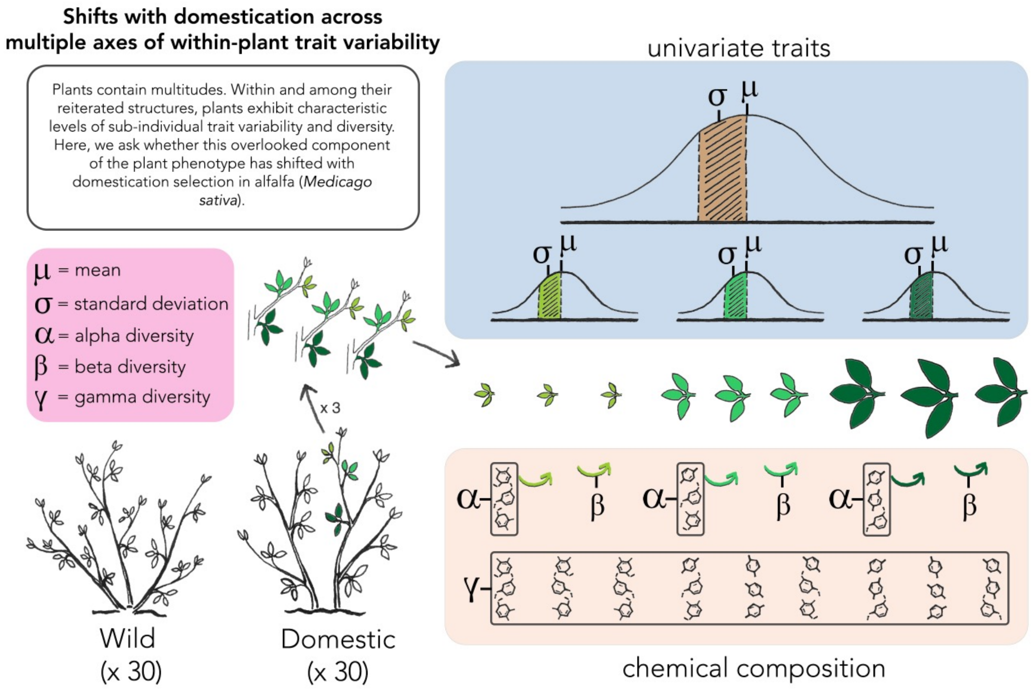

**Figure S1.2. Schematic of *Medicago sativa* domestication.** A general cartoon of alfalfa's domestication history, including some but not all of the cultivars used in this study (see Table S1.4 for a comprehensive list of the 30 wild populations and 30 cultivars studied). We used our historical understanding of alfalfa's domestication history (Barnes, 1977; Havananda *et al.*, 2010) to select wild plants representing a broad geographic extent and population/subspecies structure, and domestic cultivars to represent an array of domestication histories among those subspecies. Wild populations were selected to include three wild progenitor subspecies (*Medicago sativa* ssp. *caerulea*, ssp. *hemicycla*, and ssp. *falcata*) across a broad geographic extent of the native range. We prioritized wild accessions of known ploidy (diploid), to minimize likelihood of choosing wild populations experiencing gene flow with nearby cultivars. Commercial cultivars were selected to encompass different breeding histories within and among the wild subspecies, as well as various introductions of alfalfa germplasm into the United States. Table S1.4 provides geographic information for wild populations and backgrounds of cultivars used in this study, selected from the USDA GRIN repository.

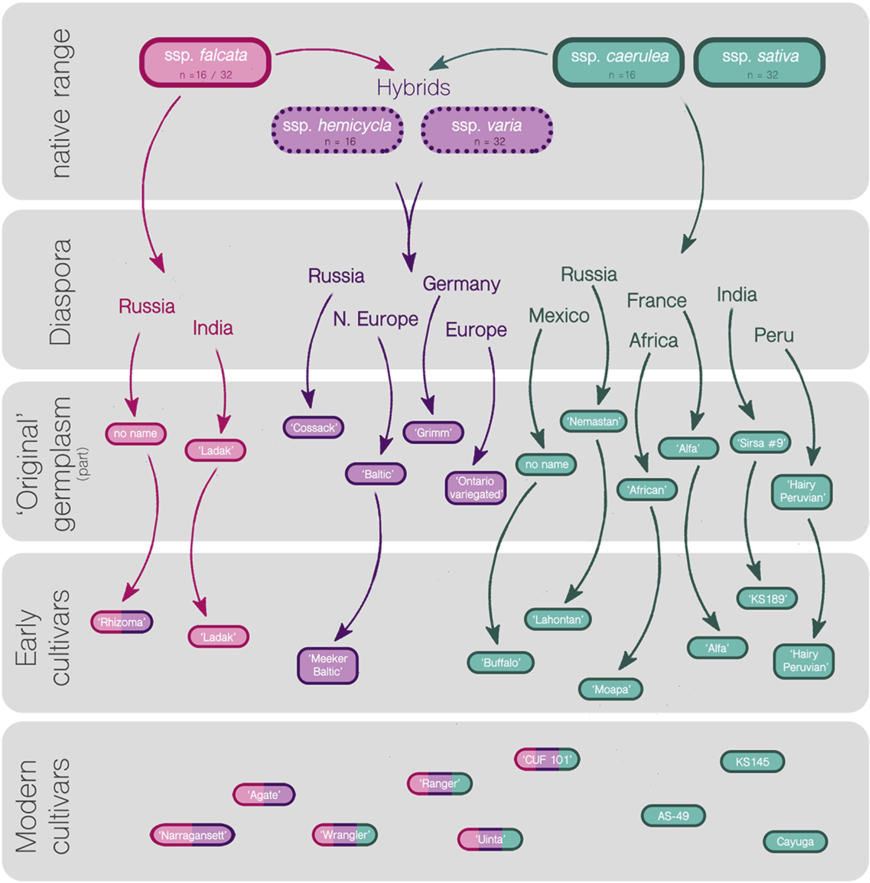

**Figure S1.3. Measures of within-plant variability for wild and domestic plants.** Variability is quantified as the standard deviation in physical traits (Specific Leaf Area, trichome density), nutritive traits (Leaf Water Content, Carbon, Nitrogen, and C:N ratio), and concentrations of N = 84 saponin compounds (data shown for of 5 random compounds). We quantified chemical richness, Shannon, and Simpson's diversity within single leaves (alpha diversity) and aggregated at the whole-plant scale (gamma diversity). We also quantified variability in terms of dissimilarity in chemical composition among leaves (beta diversity). Plots show levels of variability within leaves of young (up-triangle), middle (diamond) and older (down-triangle) age classes, as well as variability encompassed by leaves of all ontogenetic stages (circles). For chemical diversity, triangle/diamond points show the average alpha diversity of leaves within those age classes, and circles show the average gamma diversity of entire plants. For chemical dissimilarity, triangles/diamonds are the average dispersions within/among leaves of each age class, and circles show the average dispersion among all leaves, across ontogeny.

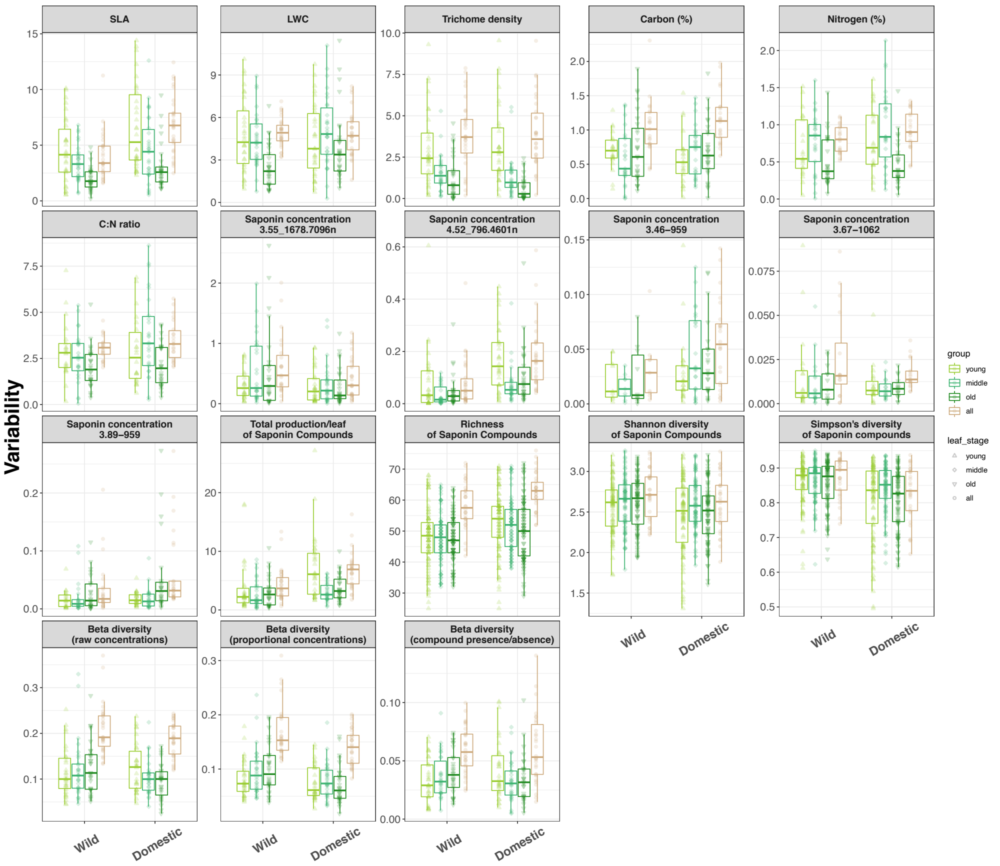

**Figure S1.4. Measures of within-plant trait means for wild and domestic plants.** Means shown correspond to the metrics of variability plotted above. For Specific Leaf Area, trichome density, Leaf Water Content, Carbon, Nitrogen, C:N ratio, concentrations of single saponin compounds, and total production of saponins, this is the trait mean that directly accompanies the trait standard deviation shown in Fig S3 (trait mean/sd among N = 3 leaves within age class, or across all N = 9 leaves/plant). Means for total compound production correspond to the covariates for compound production used in models of gamma and beta chemical diversity. For chemical richness, Shannon, and Simpson's diversity within single leaves (alpha diversity), covariates in models were the corresponding total compound concentrations per leaf (not shown).

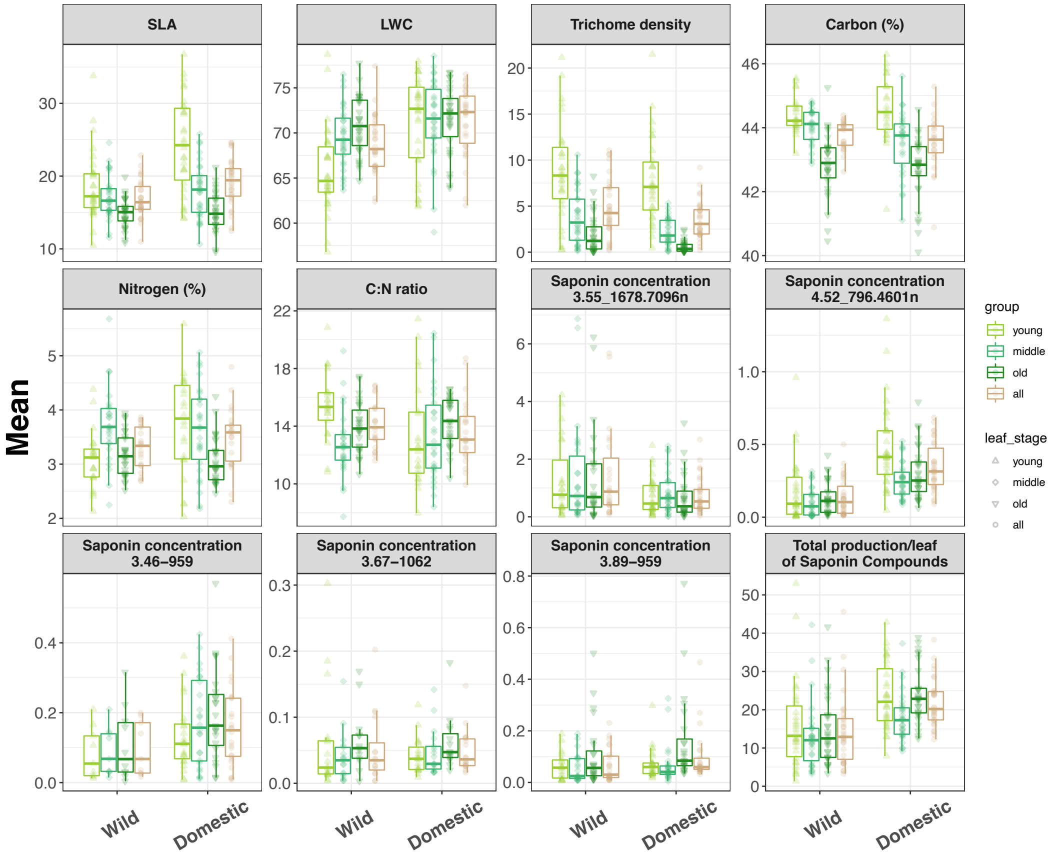

**Figure S1.5.** Size of wild and domestic plants, measured after 64 days of growth at time of harvest for plant traits. Plant size was calculated as volume (height × width1 × width2). Domestic *M. sativa* were 34.5 $cm^2$ larger, on average, compared to wild plants ($F_{(1,58)} = 18.3$, $p<0.001$).

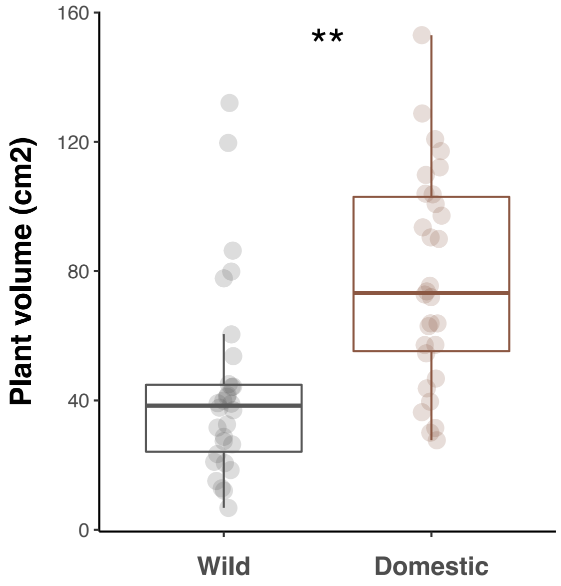

### Supplemental Tables

**Table S1.1. Effects of domestication on among-leaf trait variability in alfalfa.** Models test the hypothesis that levels of within-plant, among-leaf trait variability differ between domestic plants and their wild progenitors. Variability is quantified among all leaves per plant (whole-plant variability) as well as within leaves of different ontogenetic stages (young, middle, older age classes). Response variables are the standard deviation among leaves for physical traits (SLA, Trichome density), nutritive traits (LWC, Carbon, Nitrogen, C:N ratio) and concentrations of saponin compounds (concentration of single compounds & total concentration per leaf); the diversity of saponin compounds within individual leaves (saponin richness, Shannon diversity, and Simpson's diversity); and dissimilarity of saponin chemistry among leaves (beta diversity) in compound identity (presence/absence); proportional concentrations; and raw concentrations. In cases of significant interactions (Domestication × Leaf stage), significance is shown for post-hoc contrasts (t-tests) within leaf age class. Hyphens indicate that effects are consistent for leaves within each age class (no interactions). See Tables S2.1 and S2.2 for full models. Significance levels are indicated by asterisks: $p \leq .05$ (*), $p \leq .01$ (**), and $p \leq .001$ (***).

| Trait / Response | Leaf age class | N | *df* (residual) | β (± 95% CI) | % change (± 95% CI) | Test statistic | p-value | Sig. |
|---|---|---|---|---|---|---|---|---|
| SLA | all | 60 | 56 | 3.01 (1.75, 4.14) | 83.9% (48.9%, 115.4%) | $\chi^2 = 26.70$ | **< .001** | *** |
| SLA | young | 180 | 174 | 1.44 (0.74, 2.23) | 29.8% (15.3%, 46%) | $\chi^2 = 14.64$ | **< .001** | *** |
| SLA | middle | - | - | - | - | - | - | *** |
| SLA | old | - | - | - | - | - | - | *** |
| Trichome density | all | 60 | 57 | 0.06 (-0.99, 0.97) | 1.6% (-25.5%, 25.2%) | $\chi^2 = 0.01$ | 0.905 | |
| Trichome density | young | 180 | 174 | -0.26 (-0.72, 0.26) | -22.7% (-51.1%, 29.1%) | $\chi^2 = 1.16$ | 0.282 | |
| Trichome density | middle | - | - | - | - | - | - | |
| Trichome density | old | - | - | - | - | - | - | |
| LWC | all | 60 | 57 | -0.04 (-0.69, 0.68) | -0.8% (-14.0%, 13.7%) | $\chi^2 = 0.01$ | 0.903 | |
| LWC | young | 180 | | -0.36 (-1.41, 0.72) | -7.6% (-29.7%, 15.1%) | $t_{172} = 0.62$ | 0.534 | |
| LWC | middle | 180 | | 0.65 (-0.47, 1.81) | 14.8% (-10.7%, 41.0%) | $t_{172} = -1.12$ | 0.263 | |
| LWC | old | 180 | | 1.4 (0.25, 2.57) | 55.8% (10.0%, 102.7%) | $t_{172} = -2.41$ | **0.017** | * |
| C | all | 43 | 40 | 0.14 (-0.08, 0.34) | 14.9% (-8.2%, 35.7%) | $\chi^2 = 1.64$ | 0.201 | |
| C | young | 151 | 145 | 0.02 (-0.1, 0.16) | 3.3% (-16.6%, 26.5%) | $\chi^2 = 0.09$ | 0.767 | |

| | | | | | | | | |
|---|---|---|---|---|---|---|---|---|
| C | middle | - | - | - | - | - | - | |
| C | old | - | - | - | - | - | - | |
| N | all | 43 | 39 | 0.11 (-0.03, 0.26) | 13.6% (-3.3%, 32.1%) | $\chi^2 = 2.47$ | 0.116 | |
| N | young | 151 | 144 | 0.05 (-0.09, 0.2) | 7.4% (-12.8%, 27.3%) | $\chi^2 = 0.56$ | 0.456 | |
| N | middle | - | - | - | - | - | - | |
| N | old | - | - | - | - | - | - | |
| C:N | all | 43 | 40 | 0.41 (-0.10, 0.91) | 13.4% (-3.2%, 29.9%) | $\chi^2 = 2.34$ | 0.126 | |
| C:N | young | 151 | | -0.04 (-0.99, 0.82) | -1.5% (-34.0%, 28.2%) | $t_{144} = 0.09$ | 0.927 | |
| C:N | middle | 151 | | 1.13 (0.29, 1.98) | 43.9% (11.1%, 76.6%) | $t_{144} = -2.62$ | **0.01** | * |
| C:N | old | 151 | | 0.08 (-0.76, 0.85) | 4.1% (-37.4%, 41.8%) | $t_{144} = -0.2$ | 0.839 | |
| Saponins (single compounds) | all | 2892 | 2885 | 0.32 (0.09, 0.54) | 37.3% (9.4%, 71.3%) | $\chi^2 = 8.27$ | **0.004** | ** |
| Saponins (single compounds) | young | 8939 | | 0.42 (0.25, 0.61) | 52.2% (28.7%, 83.2%) | $t_{8929} = -4.61$ | **< .001** | *** |
| Saponins (single compounds) | middle | 8939 | | 0.35 (0.17, 0.54) | 41.8% (18.6%, 71.7%) | $t_{8929} = -3.82$ | **< .001** | *** |
| Saponins (single compounds) | old | 8939 | | 0.48 (0.29, 0.65) | 60.8% (34.1%, 90.9%) | $t_{8929} = -5.19$ | **< .001** | *** |
| Saponins (total concentration) | all | 60 | 56 | 2.1 (0.6, 3.39) | 43.3% (12.3%, 69.7.0%) | $\chi^2 = 7.83$ | **0.005** | ** |
| Saponins (total concentration) | young | 180 | | 0.66 (0.28, 1.05) | 93.3% (32.0%, 184.6%) | $t_{170} = -3.1$ | **0.002** | ** |
| Saponins (total concentration) | middle | 180 | | 0.21 (-0.23, 0.63) | 23.4% (-20.4%, 87.1%) | $t_{170} = -0.99$ | 0.325 | |
| Saponins (total concentration) | old | 180 | | 0.25 (-0.14, 0.65) | 28.1% (-13.3%, 92.4%) | $t_{170} = -1.16$ | 0.247 | |
| Richness | all ($\gamma$) | 60 | 55 | 3.91 (0.27, 7.91) | 6.6% (0.5%, 13.4%) | $\chi^2 = 3.95$ | **0.047** | * |
| Richness | young ($\alpha$) | 540 | 533 | 4.18 (2.56, 5.67) | 8.5% (5.2%, 11.5%) | $\chi^2 = 27.42$ | **<.001** | *** |
| Richness | middle ($\alpha$) | - | - | - | - | - | - | |
| Richness | old ($\alpha$) | - | - | - | - | - | - | |
| Shannon diversity | all ($\gamma$) | 60 | 55 | -0.06 (-0.24, 0.11) | -2.1% (-8.8%, 4.2%) | $\chi^2 = 0.39$ | 0.535 | |
| Shannon diversity | young ($\alpha$) | 540 | 533 | -0.04 (-0.1, 0.03) | -1.5% (-4%, 1.3%) | $\chi^2 = 1.22$ | 0.27 | |
| Shannon diversity | middle ($\alpha$) | - | - | - | - | - | - | |
| Shannon diversity | old ($\alpha$) | - | - | - | - | - | - | |
| Simpson diversity | all ($\gamma$) | 60 | 55 | -0.03 (-0.07, 0.01) | -4.0% (-8.4%, 0.7%) | $\chi^2 = 3.07$ | **0.08** | |
| Simpson diversity | young ($\alpha$) | 540 | 532 | -0.02 (-0.04, -0.01) | -3.0% (-4.6%, -1.3%) | $\chi^2 = 12.23$ | **< .001** | *** |
| Simpson diversity | middle ($\alpha$) | - | - | - | - | - | - | |

| | | | | | | | | |
|---|---|---|---|---|---|---|---|---|
| Simpson diversity | old (α) | - | - | - | - | - | - | |
| Beta diversity (presence/absence) | all | 60 | 57 | 0.00 (-0.01, 0.02) | 2.70% (19.4%, 22.5%) | $\chi^2 = 0.06$ | 0.807 | |
| Beta diversity (presence/absence) | young | 180 | | 0.26 (-0.06, 0.58) | 30.2% (-6%, 77.8%) | $t_{172} = -1.71$ | 0.089 | |
| Beta diversity (presence/absence) | middle | 180 | | -0.17 (-0.46, 0.12) | -16.0% (-37.1%, 12.8%) | $t_{172} = 1.13$ | 0.259 | |
| Beta diversity (presence/absence) | old | 180 | | -0.21 (-0.5, 0.08) | -19.1% (-39.1%, 8.3%) | $t_{172} = 1.37$ | 0.171 | |
| Beta diversity (proportional) | all | 60 | 56 | -0.15 (-0.27, -0.02) | -13.7% (-23.3%, -2.0%) | $\chi^2 = 5.35$ | **0.021** | * |
| Beta diversity (proportional) | young | 180 | 174 | -0.22 (-0.35, -0.08) | -19.4% (-29.5%, -7.5%) | $\chi^2 = 9.41$ | **0.002** | ** |
| Beta diversity (proportional) | middle | - | - | - | - | - | - | |
| Beta diversity (proportional) | old | - | - | - | - | - | - | |
| Beta diversity (raw) | all | 60 | 56 | -0.02 (-0.04, 0.01) | -8.0 % (-20.3%, 4.0%) | $\chi^2 = 1.92$ | 0.166 | |
| Beta diversity (raw) | young | 180 | | 0.1 (-0.12, 0.32) | 10.4 % (-11.4%, 38.2%) | $t_{172} = -0.89$ | 0.374 | |
| Beta diversity (raw) | middle | 180 | | -0.1 (-0.31, 0.12) | -9.3 % (-26.7%, 13.1%) | $t_{172} = 0.88$ | 0.382 | |
| Beta diversity (raw) | old | 180 | | -0.28 (-0.51, -0.05) | -24.1 % (-39.7%, -4.5%) | $t_{172} = 2.47$ | **0.014** | * |

**Table S1.2. Effects of domestication on among-leaf trait means in alfalfa.** Models test the hypothesis that traits means differ between domestic plants and their wild progenitors. Trait means are quantified among all leaves per plant (plant-level average) as well as across leaf ontogeny (within three leaf age classes). Response variables are the among-leaf average for physical traits (SLA, Trichome density), nutritive traits (LWC, Carbon, Nitrogen, C:N ratio) and concentrations of saponin compounds (concentration of single compounds & total concentration per leaf). In cases of significant interactions (Domestication × Leaf stage), significance is shown for post-hoc contrasts (t-tests) within leaf age class. Hyphens indicate that effects are consistent for leaves within each age class (no interactions). See Tables S2.1 and S2.2 for full models. Significance levels are indicated by asterisks: $p \leq .05$ (*), $p \leq .01$ (**), and $p \leq .001$ (***).

| Trait / Response | Leaf age class | N | *df* (residual) | β (± 95% CI) | % change (± 95% CI) | Test statistic | p-value | Sig. |
|---|---|---|---|---|---|---|---|---|
| SLA | all | 60 | 57 | 2.29 (0.78, 4.03) | 13.5% (4.6%, 23.9%) | $\chi^2 = 9.3$ | **0.002** | ** |
| SLA | young | 180 | | 5.68 (3.71, 7.92) | 30.5% (20%, 42.6%) | $t_{172} = -5.5$ | **< .001** | *** |
| SLA | middle | 180 | | 0.94 (-1.15, 3.25) | 5.5% (-6.7%, 19.0%) | $t_{172} = -0.92$ | 0.361 | |
| SLA | old | 180 | | 0.23 (-1.8, 2.12) | 1.6% (-12.1%, 14.2%) | $t_{172} = -0.23$ | 0.821 | |
| Trichome density | all | 60 | 57 | -1.33 (-2.54, -0.03) | -27.4% (-52.4%, -0.5%) | $\chi^2 = 3.81$ | **0.051** | * |
| Trichome density | young | 180 | 174 | -1.33 (-2.67, 0.19) | -14.9% (-29.9%, 2.1%) | $\chi^2 = 3.82$ | **0.051** | * |
| Trichome density | middle | - | - | - | - | - | - | * |
| Trichome density | old | - | - | - | - | - | - | * |
| LWC | all | 60 | 57 | 2.61 (0.79, 4.58) | 3.8% (1.1%, 6.7%) | $\chi^2 = 8.68$ | **0.003** | ** |
| LWC | young | 180 | | 5.45 (3.48, 7.65) | 8.3% (5.3%, 11.6%) | $t_{172} = -5.14$ | **< .001** | *** |
| LWC | middle | 180 | | 1.78 (-0.18, 3.99) | 2.6% (-0.3%, 5.7%) | $t_{172} = -1.68$ | 0.095 | |
| LWC | old | 180 | | 0.61 (-1.48, 2.74) | 0.9% (-2.1%, 3.9%) | $t_{172} = -0.57$ | 0.567 | |
| C | all | 43 | 40 | -0.21 (-0.61, 0.23) | -0.5% (-1.4%, 0.5%) | $\chi^2 = 0.62$ | 0.433 | |
| C | young | 151 | | 0.18 (-0.35, 0.75) | 0.4% (-0.8%, 1.7%) | $t_{143} = -0.7$ | 0.486 | |
| C | middle | 151 | | -0.47 (-0.97, 0.03) | -1.1% (-2.2%, 0.1%) | $t_{143} = 1.88$ | 0.062 | |
| C | old | 151 | | -0.17 (-0.63, 0.34) | -0.4% (-1.5%, 0.8%) | $t_{143} = 0.71$ | 0.478 | |
| N | all | 43 | 39 | 0.24 (-0.05, 0.49) | 7.7% (-1.7%, 15.6%) | $\chi^2 = 3.05$ | 0.081 | |
| N | young | 151 | | 0.69 (0.31, 1.1 ) | 23.3% (10.3%, 36.9%) | $t_{143} = -3.53$ | **0.001** | *** |

| | | | | | | | | |
|---|---|---|---|---|---|---|---|---|
| N | middle | 151 | | -0.02 (-0.34, 0.36) | -0.5% (-9.4%, 10.0%) | $t_{143}$ = 0.09 | 0.93 | |
| N | old | 151 | | -0.17 (-0.63, 0.34) | -0.4% (-1.5%, 0.8%) | $t_{143}$ = 0.71 | 0.478 | |
| C:N | all | 43 | 39 | -0.67 (-1.69, 0.47) | -4.5% (-11.5%, 3.2%) | $\chi^2$ = 1.43 | 0.232 | |
| C:N | young | 151 | | -2.07 ( -3.54, -0.39) | -13.2% (-22.6%, -2.5%) | $t_{142}$ = 2.79 | **0.006** | ** |
| C:N | middle | 151 | | 0.57 (-0.95, 1.97) | 4.3% (-7.2%, 14.9%) | $t_{142}$ = -0.83 | 0.406 | |
| C:N | old | 151 | | 0.28 (-1.01, 1.6) | 1.9% (-7.0%, 11.1%) | $t_{142}$ = -0.42 | 0.674 | |
| Saponins (single compounds) | all | 2892 | 2885 | 0.37 (0.18, 0.57) | 44.4% (20.0%, 76%) | $\chi^2$ = 14.8 | **< .001** | *** |
| Saponins (single compounds) | young | 8939 | 8931 | 0.42 (0.27, 0.59) | 52.2% (30.8%, 80.1%) | $\chi^2$ = 26.78 | **< .001** | *** |
| Saponins (single compounds) | middle | - | - | - | - | - | - | *** |
| Saponins (single compounds) | old | - | - | - | - | - | - | *** |
| Saponins (total concentration) | all | 60 | 55 | 6.04 (1.65, 10.32) | 38.0% (10.4%, 64.9%) | $\chi^2$ = 6.64 | **0.01** | ** |
| Saponins (total concentration) | young | 180 | | 0.57 (0.29, 0.86) | 76.0% (33.9%, 136.3%) | $t_{170}$ = -3.56 | **< .001** | *** |
| Saponins (total concentration) | middle | 180 | | 0.43 (0.16, 0.74) | 54.1% (16.8%, 109.4%) | $t_{170}$ = -2.72 | **0.007** | ** |
| Saponins (total concentration) | old | 180 | | 0.57 (0.24, 0.85) | 77.0% (27.2%, 134.6%) | $t_{170}$ = -3.6 | **<.001** | *** |

**Table S1.3. Effects of domestication on within-plant variability, accounting for shifts in associated means or total compound production.** Models test the hypothesis that differences within-plant trait variability between domestic plants and their wild progenitors are explained by shifts in associated trait means, or shifts in overall production of secondary metabolites – e.g. the relationship between trait variability (Table S1) and trait means or magnitudes (Table S2). Response variables are the standard deviation among leaves for physical traits (SLA, Trichome density), nutritive traits (LWC, Carbon, Nitrogen, C:N ratio) and concentrations of saponin compounds (concentration of single compounds & total concentration per leaf); the diversity of saponin compounds within individual leaves (saponin richness, Shannon diversity, and Simpson's diversity); and dissimilarity of saponin chemistry among leaves (beta diversity) in compound identity (presence/absence); proportional concentrations; and raw concentrations. For chemical diversity, we used the peak area at the corresponding scale as a covariate to parallel use of the mean for other traits (total concentration per leaf for α diversity; and averaged across all leaves or within leaf age class for $\gamma$ and β diversity at the whole-plant and within age-class scales, respectively). In cases of significant interactions (Domestication × Leaf stage or Domestication × Mean), significance is shown for post-hoc contrasts (t-tests) within leaf age class and/or holding the mean constant at its average value. Hyphens indicate that effects are consistent for leaves within each age class (no interactions). See Tables S2.1 and S2.2 for full models. Significance levels are indicated by asterisks: $p \leq .05$ (*), $p \leq .01$ (**), and $p \leq .001$ (***).

| Trait / Response | Leaf age class | N | *df* (residual) | β (± 95% CI) | % change (± 95% CI) | Test statistic | p-value | Sig. |
|---|---|---|---|---|---|---|---|---|
| SLA | all | 60 | 55 | 1.71 (0.85, 2.61) | 39.3% (19.7%, 60.1%) | $\chi^2 = 15.0$ | **< .001** | *** |
| SLA | young | 180 | 173 | 0.79 (0.18, 1.45) | 18.8% (4.2%, 34.5%) | $\chi^2 = 5.25$ | **0.022** | ** |
| SLA | middle | - | - | - | - | - | - | ** |
| SLA | old | - | - | - | - | - | - | ** |
| Trichome density | all | 60 | - | 0.37 (0.11, 0.62) | 45.0 % (11.8%, 85.8%) | $t_{55} = -2.81$ | **0.007** | ** |
| Trichome density | young | 180 | 173 | 0.16 (-0.06, 0.37) | 17.2% (-5.6%, 44.3%) | $\chi^2 = 1.95$ | 0.163 | |
| Trichome density | middle | - | - | - | - | - | - | |
| Trichome density | old | - | - | - | - | - | - | |
| LWC | all | 60 | 56 | 0.35 (-0.26, 0.98) | 7.5% (-5.6%, 20.7%) | $\chi^2 = 1.19$ | 0.274 | |
| LWC | young | 180 | 173 | 0.93 (0.24, 1.55) | 23.9% (6.1%, 39.8%) | $\chi^2 = 7.28$ | **0.007** | ** |
| LWC | middle | - | - | - | - | - | - | ** |

| | | | | | | | | |
|---|---|---|---|---|---|---|---|---|
| LWC | old | - | - | - | - | - | - | ** |
| C | all | 43 | 38 | 0.13 (-0.1, 0.36) | 13.6% (-10.3%, 37.5%) | $\chi^2 = 1.36$ | 0.244 | |
| C | young | 151 | 142 | -0.04 (-0.18, 0.12) | -6.0% (-27.6%, 18.6%) | $\chi^2 = 0.32$ | 0.573 | |
| C | middle | - | - | - | - | - | - | |
| C | old | - | - | - | - | - | - | |
| N | all | 43 | 38 | 0.06 (-0.06, 0.18) | 6.8% (-7.1%, 21.2%) | $\chi^2 = 0.92$ | 0.338 | |
| N | young | 151 | 143 | 0.02 (-0.1, 0.14) | 2.8% (-14.0%, 19.0%) | $\chi^2 = 0.09$ | 0.764 | |
| N | middle | - | - | - | - | - | - | |
| N | old | - | - | - | - | - | - | |
| C:N | all | 43 | | 0.51 (0.06, 0.95) | 16.8% (2.1%, 31.3%) | $t_{38} = -2.21$ | **0.033** | * |
| C:N | young | 151 | | -0.05 (-0.88, 0.97) | -1.5% (-29.0%, 31.9%) | $t_{137} = 0.09$ | 0.925 | |
| C:N | middle | 151 | | 1.23 (0.43, 1.98) | 46.7% (16.5%, 75.2%) | $t_{137} = -2.94$ | **0.004** | ** |
| C:N | old | 151 | | 0.04 (-0.74, 0.81) | 2.0% (-36.7%, 40.5%) | $t_{137} = -0.11$ | 0.915 | |
| Saponins (single compounds) | all | 2892 | 2880 | -0.01 (-0.12, 0.09) | -1.4% (-11.0%, 9.5%) | $\chi^2 = 0.07$ | 0.786 | |
| Saponins (single compounds) | young | 8939 | | 0.04 (-0.06, 0.14) | 3.6% (-6.2%, 15.0%) | $t_{8923} = -0.71$ | 0.479 | |
| Saponins (single compounds) | middle | 8939 | | -0.05 (-0.15, 0.06) | -4.6% (-14.0%, 5.8%) | $t_{8923} = 0.94$ | 0.345 | |
| Saponins (single compounds) | old | 8939 | | -0.07 (-0.18, 0.04) | -6.6% (-16.3%, 3.8%) | $t_{8923} = 1.33$ | 0.182 | |
| Saponins (total concentration) | all | 60 | 55 | 0.82 (-0.64, 2.18) | 15.4% (-11.9%, 40.5%) | $\chi^2 = 1.39$ | 0.238 | |
| Saponins (total concentration) | young | 180 | 171 | 0.13 (-0.16, 0.4) | 13.7% (-14.4%, 48.5%) | $\chi^2 = 0.88$ | 0.349 | |
| Saponins (total concentration) | middle | - | - | - | - | - | - | |
| Saponins (total concentration) | old | - | - | - | - | - | - | |
| Richness | all ($\gamma$) | 60 | 54 | -0.69 (-4.41, 2.46) | -1.1% (-7.2%, 4.0%) | $\chi^2 = 0.18$ | 0.674 | |
| Richness | young ($\alpha$) | 540 | | -0.21 (-3.48, 3.12) | -0.4% (-7.0%, 6.3%) | $t_{526} = 0.12$ | 0.904 | |
| Richness | middle ($\alpha$) | 540 | | 0.70 (-2.66, 3.90) | 1.4% (-5.3%, 7.8 %) | $t_{526} = -0.41$ | 0.685 | |
| Richness | old ($\alpha$) | 540 | | -1.45 (-5.10, 2.03) | -3.0% (-10.5%, 4.20%) | $t_{526} = 0.80$ | 0.423 | |
| Shannon diversity | all ($\gamma$) | 60 | 54 | -0.06 (-0.26, 0.13) | -2.2% (-9.4%, 4.7%) | $\chi^2 = 0.38$ | 0.538 | |
| Shannon diversity | young ($\alpha$) | 540 | | -0.05 (-0.23, 0.12) | -1.8% (-9.1%, 4.8%) | $t_{523} = 0.55$ | 0.582 | |
| Shannon diversity | middle ($\alpha$) | 540 | | 0.10 (-0.08, 0.27) | 3.8% (-3.0%, 10.0%) | $t_{523} = -1.20$ | 0.232 | |
| Shannon diversity | old ($\alpha$) | 540 | | -0.00 (-0.16, 0.18) | -0.2% (-6.3%, 6.9%) | $t_{523} = -0.05$ | 0.962 | |

| | | | | | | | | |
|---|---|---|---|---|---|---|---|---|
| Simpson diversity | all ($\gamma$) | 60 | 53 | -0.04 (-0.08, -0.00) | -4.5% (-9.1%, 0.5%) | $\chi^2 = 3.56$ | 0.059 | |
| Simpson diversity | young (α) | 540 | | -0.05 (-0.08, -0.01) | -5.5% (-9.4%, -1.0%) | $t_{523} = 2.45$ | **0.015** | * |
| Simpson diversity | middle (α) | 540 | | -0.00 (-0.04, 0.03) | -0.5% (-4.6%, 4.0%) | $t_{523} = 0.25$ | 0.805 | |
| Simpson diversity | old (α) | 540 | | -0.02 (-0.06, 0.02) | -2.6% (-7.1%, 2.2%) | $t_{523} = 1.13$ | 0.260 | |
| Beta diversity (presence/absence) | all | 60 | 56 | 0.01 (0.00, 0.02) | 18.1% (-3.9%, 38.1%) | $\chi^2 = 2.29$ | 0.130 | |
| Beta diversity (presence/absence) | young | 180 | | 0.36 (0.06, 0.69) | 43.1% (5.7%, 98.5%) | $t_{171} = -2.31$ | **0.022** | * |
| Beta diversity (presence/absence) | middle | 180 | | -0.11 (-0.39, 0.21) | -10.6% (-32.3%, 23.3%) | $t_{171} = 0.74$ | 0.459 | |
| Beta diversity (presence/absence) | old | 180 | | -0.10 (-0.40, 0.20) | -9.7% (-32.7%, 21.8%) | $t_{171} = 0.65$ | 0.518 | |
| Beta diversity (proportional) | all | 60 | 56 | -0.15 (-0.28, -0.02) | -13.7% (-24.0%, -2.1%) | $\chi^2 = 4.61$ | **0.032** | * |
| Beta diversity (proportional) | young | 180 | 173 | -0.18 (-0.33, -0.02) | -16.1% (-28.2%, -1.8%) | $\chi^2 = 5.67$ | **0.017** | * |
| Beta diversity (proportional) | middle | - | - | - | - | - | - | * |
| Beta diversity (proportional) | old | - | - | - | - | - | - | * |
| Beta diversity (raw) | all | 60 | 55 | -0.00 (-0.03, 0.02) | -2.0% (-13.7%, 9.1%) | $\chi^2 = 0.11$ | 0.743 | |
| Beta diversity (raw) | young | 180 | | 0.14 (-0.07, 0.37) | 15.3% (-6.9%, 45.0%) | $t_{170} = -1.30$ | 0.196 | |
| Beta diversity (raw) | middle | 180 | | -0.08 (-0.3, 0.13) | -8.1% (-25.7%, 14.1%) | $t_{170} = 0.79$ | 0.432 | |
| Beta diversity (raw) | old | 180 | | -0.22 (-0.44, 0.01) | -19.6% (-35.4%, 0.6%) | $t_{170} = 1.96$ | **0.052** | * |

**Table S1.4.** Plant populations and cultivars used in this study. Wild populations were selected to encompass multiple progenitor subspecies, across each subspecies' geographic range (Fig S2). Domestic cultivars were selected to include a variety of domestication histories, both pre- and post-arrival into the United States (Barnes, 1977) (Table S5). Estimated percents of each wild subspecies were estimated using Barnes (1977), which provides percentage breakdowns of each cultivar according to wild subspecies or among each of 6 single-source introductions (each of which can, in turn, be traced to one of the three wild subspecies). See Barnes (1977), Table 2.

| Population / cultivar ID | GRIN Accession ID | Domestication status | wild ssp or cultivar name | Source population* | Estimated % of each wild ssp | | |
|---|---|---|---|---|---|---|---|
| | | | | | ssp. *falcata* | ssp. *hemicycla* | ssp. *caerulea* |
| 01_W_FAL | PI 464729 | W | ssp. falcata | Turkey | 100 | 0 | 0 |
| 02_W_FAL | PI 577555 | W | ssp. falcata | Ukraine | 100 | 0 | 0 |
| 03_W_FAL | PI 634106 | W | ssp. falcata | Ukraine | 100 | 0 | 0 |
| 04_W_FAL | PI 631811 | W | ssp. falcata | Kazakhstan | 100 | 0 | 0 |
| 05_W_FAL | PI 641544 | W | ssp. falcata | Mongolia | 100 | 0 | 0 |
| 06_W_FAL | PI 494662 | W | ssp. falcata | Romania | 100 | 0 | 0 |
| 07_W_FAL | PI 577558 | W | ssp. falcata | Russia | 100 | 0 | 0 |
| 08_W_FAL | PI 258752 | W | ssp. falcata | Russia | 100 | 0 | 0 |
| 09_W_FAL | PI 325387 | W | ssp. falcata | Russia | 100 | 0 | 0 |
| 10_W_FAL | PI 631707 | W | ssp. falcata | China | 100 | 0 | 0 |
| 11_W_HEM | PI 631814 | W | ssp. hemicycla | Russia | 0 | 100 | 0 |
| 12_W_HEM | PI 315460 | W | ssp. hemicycla | Russia | 0 | 100 | 0 |
| 13_W_HEM | PI 464727 | W | ssp. hemicycla | Turkey | 0 | 100 | 0 |
| 14_W_HEM | PI 464728 | W | ssp. hemicycla | Turkey | 0 | 100 | 0 |
| 15_W_HEM | PI 577543 | W | ssp. hemicycla | Georgia | 0 | 100 | 0 |
| 16_W_HEM | PI 634176 | W | ssp. hemicycla | Kazakhstan | 0 | 100 | 0 |
| 17_W_HEM | PI 634136 | W | ssp. hemicycla | Kazakhstan | 0 | 100 | 0 |
| 18_W_HEM | PI 641615 | W | ssp. hemicycla | Kazakhstan | 0 | 100 | 0 |
| 19_W_HEM | PI 641619 | W | ssp. hemicycla | Kazakhstan | 0 | 100 | 0 |
| 20_W_HEM | PI 641601 | W | ssp. hemicycla | Kazakhstan | 0 | 100 | 0 |
| 21_W_CAER | PI 577549 | W | ssp. caerulea | Georgia | 0 | 0 | 100 |
| 22_W_CAER | PI 440500 | W | ssp. caerulea | Kazakhstan | 0 | 0 | 100 |
| 23_W_CAER | PI 641606 | W | ssp. caerulea | Kazakhstan | 0 | 0 | 100 |
| 24_W_CAER | PI 577541 | W | ssp. caerulea | Kazakhstan | 0 | 0 | 100 |

| | | | | | | | |
|---|---|---|---|---|---|---|---|
| 25_W_CAER | PI 464713 | W | ssp. caerulea | Turkey | 0 | 0 | 100 |
| 26_W_CAER | PI 464721 | W | ssp. caerulea | Turkey | 0 | 0 | 100 |
| 27_W_CAER | PI 464723 | W | ssp. caerulea | Turkey | 0 | 0 | 100 |
| 28_W_CAER | PI 210367 | W | ssp. caerulea | Iran | 0 | 0 | 100 |
| 29_W_CAER | PI 314267 | W | ssp. caerulea | Uzbekistan | 0 | 0 | 100 |
| 30_W_CAER | PI 505871 | W | ssp. caerulea | Former Soviet Union | 0 | 0 | 100 |
| 31_D_RHI | W6 2525 | D | Rhizoma | Europe | 50 | 50 | 0 |
| 32_D_LAD | W6 2502 | D | Ladak | India | 90 | 5 | 5 |
| 33_D_MEE | PI 672756 | D | Meeker Baltic | Europe | 0 | 100 | 0 |
| 34_D_LAH | W6 2516 | D | Lahontan | Russia, Middle East | 5 | 5 | 90 |
| 35_D_ALF | W6 2506 | D | Alfa | France | 0 | 0 | 100 |
| 36_D_BUF | W6 2498 | D | Buffalo | Mexico / South America | 0 | 0 | 100 |
| 37_D_HAI | PI 672744 | D | Hairy Peruvian | Peru | 0 | 0 | 100 |
| 38_D_KS189 | PI 672752 | D | KS189 | India | 0 | 0 | 100 |
| 39_D_MOA | W6 2518 | D | Moapa | North Africa | 0 | 0 | 100 |
| 40_D_KS145 | PI 672750 | D | KS145 | mix | 0 | 0 | 100 |
| 41_D_UIN | PI 672766 | D | Uinta | mix | 12 | 51 | 37 |
| 42_D_AGA | W6 2505 | D | Agate | mix | 47 | 40 | 13 |
| 43_D_NAR | W6 2519 | D | Narragansett | mix | 25 | 75 | 0 |
| 44_D_WRA | PI 601132 | D | Wrangler | mix | 10 | 20 | 70 |
| 45_D_AS49 | PI 672732 | D | AS-49 | mix | 0 | 0 | 100 |
| 46_D_CUF | PI 517240 | D | CUF 101 | mix | 1 | 2 | 97 |
| 47_D_CAY | W6 2511 | D | Cayuga | mix | 11 | 15 | 74 |
| 48_D_SAR | W6 2526 | D | Saranac AR | mix | 2 | 7 | 91 |
| 49_D_WIN | PI 608671 | D | Winema | mix | 33 | 33 | 33 |
| 50_D_ATL | PI 672733 | D | Atlantic | mix | 15 | 50 | 35 |
| 51_D_RAN | PI 612887 | D | Ranger | mix | 10 | 45 | 45 |
| 52_D_VER | NSL 4099 | D | Vernal | mix | 33 | 50 | 17 |
| 53_D_ZIA | PI 672767 | D | Zia 16-2 | mix | 0 | 0 | 100 |
| 54_D_TET | PI 672768 | D | Teton | mix | 50 | 0 | 50 |
| 55_D_FLO | PI 672743 | D | Florida 77 | mix | 33 | 33 | 33 |
| 56_D_ARC | W6 2507 | D | Arc | mix | 6 | 31 | 63 |
| 57_D_MAR | W6 2517 | D | Mark II | mix | 25 | 75 | 0 |
| 58_D_GUA | PI 639220 | D | Guardsman II | mix | 33 | 33 | 33 |

| | | | | | | | |
|---|---|---|---|---|---|---|---|
| 59_D_CAL | PI 672735 | D | Caliente | mix | 0 | 0 | 100 |
| 60_D_WL202 | PI 672776 | D | WL 202 | mix | 33 | 51 | 16 |

* For wild plants, this column indicates location (country) of the wild population. For domestic plants, country names indicate where plants were cultivated before being brought to the United States. These original nine germplasm 'sources' therefore represent unique domestication histories, in different parts of the world. The remaining 21 domestic cultivars ("mix") were created by hybridizing among these original nine cultivars. Percent contributions of each wild subspecies to each cultivar were estimated using Barnes (1977).

**Table S1.5.** List of N=86 saponin compounds used in this study. See supplemental methods and materials for details of compound identification.

| ID | Integration method | m/z | Adducts | Predicted formula | Neutral mass | RT | RMD |
|---|---|---|---|---|---|---|---|
| 1 | QL | 943.5255 | [M+H]+ | C48H78O18 | 942.5177 | 4.39 | 556.95368 |
| 2 | PG | 1121.5739 | [M+H]+ | C54H88O24 | 1120.5661 | 3.43 | 508.83997 |
| 3 | QL | 959.4839 | [M+H]+ | C47H74O20 | 958.4761 | 3.46 | 504.33363 |
| 4 | PG | 1564.7015 | [M+NH4]+ | C69H110O38 | 1546.6675 | 3.56 | 448.32832 |
| 5 | PG | 1696.7433 | [M+NH4]+ | C74H118O42 | 1678.7093 | 3.56 | 438.07452 |
| 6 | PG | 1680.7473 | [M+NH4]+ | C74H118O41 | 1662.7133 | 3.58 | 444.62365 |
| 7 | PG | 1386.6191 | [M+NH4]+ | C62H96O33 | 1368.5851 | 3.59 | 446.40958 |
| 8 | PG | 1254.575 | [M+NH4]+ | C57H88O29 | 1236.541 | 3.59 | 458.32254 |
| 9 | PG | 1432.6592 | [M+NH4]+ | C64H102O34 | 1414.6252 | 3.6 | 460.12338 |
| 10 | PG | 989.4944 | [M+H]+ | C48H76O21 | 988.4866 | 3.64 | 499.64911 |
| 11 | QL | 1122.533 | [M+NH4]+ | C52H80O25 | 1104.499 | 3.64 | 474.819 |
| 12 | PG | 1240.5955 | [M+NH4]+ | C57H90O28 | 1222.5615 | 3.65 | 480.01141 |
| 13 | PG | 850.5154 | [M+NH4]+ | C42H72O16 | 832.4814 | 3.66 | 605.9855 |
| 14 | PG | 1372.6372 | [M+NH4]+ | C62H98O32 | 1354.6032 | 3.668 | 464.21589 |
| 15 | QL | 843.4363 | [M+H]+ | C42H66O17 | 842.4285 | 3.67 | 517.28862 |
| 16 | PG | 1108.552 | [M+NH4]+ | C52H82O24 | 1090.518 | 3.68 | 497.94687 |
| 17 | PG | 700.4287 | [M+NH4]+ | C36H58O12 | 682.3947 | 3.69 | 612.05373 |
| 18 | PG | 1680.7478 | [M+NH4]+ | C74H118O41 | 1662.7138 | 3.67 | 444.921 |
| 19 | PG | 1370.6246 | [M+NH4]+ | C62H96O32 | 1352.5906 | 3.73 | 455.70465 |
| 20 | PG | 1284.6232 | [M+NH4]+ | C59H94O29 | 1266.5892 | 3.71 | 485.1228 |
| 21 | QL | 1122.5686 | [M+NH4]+ | C53H84O24 | 1104.5346 | 3.79 | 506.51693 |
| 22 | QL | 1386.6535 | [M+NH4]+ | C63H100O32 | 1368.6195 | 3.76 | 471.27851 |
| 23 | QL | 1106.5372 | [M+NH4]+ | C52H80O24 | 1088.5032 | 3.79 | 485.47848 |
| 24 | PG | 1002.49 | [M+NH4]+ | C48H72O21 | 984.456 | 3.8 | 488.58352 |
| 25 | QL | 1092.5579 | [M+NH4]+ | C52H82O23 | 1074.5239 | 3.83 | 510.63655 |
| 26 | PG | 834.5207 | [M+NH4]+ | C42H72O15 | 816.4867 | 3.85 | 623.95097 |
| 27 | PG | 846.4849 | [M+NH4]+ | C42H68O16 | 828.4509 | 3.9 | 572.83952 |
| 28 | PG | 971.4843 | [M+H]+ | C48H74O20 | 970.4765 | 3.95 | 498.51552 |
| 29 | PG | 1076.5639 | [M+NH4]+ | C52H82O22 | 1058.5299 | 3.95 | 521.93933 |
| 30 | QL | 825.4274 | [M+H]+ | C42H64O16 | 824.4196 | 4 | 517.79236 |

| | | | | | | | |
|---|---|---|---|---|---|---|---|
| 31 | PG | 846.4849 | [M+NH4]+ | C42H68O16 | 828.4509 | 4.02 | 572.83952 |
| 32 | PG | 930.5058 | [M+NH4]+ | C46H72O18 | 912.4718 | 4.18 | 536.2714 |
| 33 | PG | 672.4684 | [M+NH4]+ | C36H62O10 | 654.4344 | 4.26 | 696.53831 |
| 34 | PG | 930.5418 | [M+NH4]+ | C47H76O17 | 912.5078 | 4.36 | 582.24144 |
| 35 | PG | 1059.567 | [M+H]+ | C53H86O21 | 1058.5592 | 4.4 | 535.12425 |
| 36 | PG | 652.4415 | [M+NH4]+ | C36H58O9 | 634.4075 | 4.36 | 676.68902 |
| 37 | PG | 1029.5268 | [M+H]+ | C51H80O21 | 1028.519 | 4.48 | 511.69139 |
| 38 | PG | 830.4896 | [M+NH4]+ | C42H68O15 | 812.4556 | 4.5 | 589.53177 |
| 39 | PG | 797.4678 | [M+H]+ | C42H68O14 | 796.46 | 4.53 | 586.60676 |
| 40 | PG | 816.5107 | [M+NH4]+ | C42H70O14 | 798.4767 | 4.56 | 625.46639 |
| 41 | PG | 670.4519 | [M+NH4]+ | C36H60O10 | 652.4179 | 4.59 | 674.023 |
| 42 | PG | 941.5033 | [M+H]+ | C48H76O18 | 940.4955 | 4.7 | 534.57062 |
| 43 | PG | 1056.5736 | [M+NH4]+ | C53H82O20 | 1038.5396 | 4.72 | 542.88693 |
| 44 | PG | 1069.5588 | [M+H]+ | C54H84O21 | 1068.551 | 4.74 | 522.45842 |
| 45 | PG | 652.4419 | [M+NH4]+ | C36H58O9 | 634.4079 | 4.97 | 677.30169 |
| 46 | PG | 654.4585 | [M+NH4]+ | C36H60O9 | 636.4245 | 5.3 | 700.57918 |
| 47 | PG | 1238.5797 | [M+NH4]+ | C57H88O28 | 1220.5457 | 3.76 | 469.40799 |
| 48 | QL | 1105.5431 | [M+H]+ | C53H84O24 | 1104.5353 | 3.44 | 491.25177 |
| 49 | QL | 1091.5636 | [M+H]+ | C53H86O23 | 1090.5558 | 3.49 | 516.32356 |
| 50 | QL | 1105.5789 | [M+H]+ | C54H88O23 | 1104.5711 | 3.54 | 523.61708 |
| 51 | QL | 1324.5802 | [M+NH4]+ | C60H90O31 | 1306.5462 | 3.69 | 438.02557 |
| 52 | QL | 1061.5161 | [M+H]+ | C51H80O23 | 1060.5083 | 3.69 | 486.1914 |
| 53 | QL | 1416.6571 | [M+NH4]+ | C64H102O33 | 1398.6231 | 3.7 | 463.83843 |
| 54 | QL | 1548.707 | [M+NH4]+ | C69H110O37 | 1530.673 | 3.71 | 456.50985 |
| 55 | QL | 1254.6108 | [M+NH4]+ | C58H92O28 | 1236.5768 | 3.73 | 486.84421 |
| 56 | QL | 973.501 | [M+H]+ | C48H76O20 | 972.4932 | 3.75 | 514.63738 |
| 57 | QL | 1254.6107 | [M+NH4]+ | C58H92O28 | 1236.5767 | 3.79 | 487.87988 |
| 58 | QL | 944.448 | [M+NH4]+ | C45H66O20 | 926.414 | 3.81 | 474.35116 |
| 59 | QL | 960.5162 | [M+NH4]+ | C47H74O19 | 942.4822 | 3.86 | 537.41936 |
| 60 | QL | 1090.5428 | [M+NH4]+ | C52H80O23 | 1072.5088 | 3.87 | 497.73379 |
| 61 | QL | 832.505 | [M+NH4]+ | C42H70O15 | 814.471 | 3.95 | 606.60296 |
| 62 | QL | 1087.568 | [M+H]+ | C54H86O22 | 1086.5602 | 4.06 | 513.25989 |
| 63 | QL | 832.505 | [M+NH4]+ | C42H70O15 | 814.471 | 4.07 | 606.60296 |
| 64 | QL | 959.52 | [M+H]+ | C48H78O19 | 958.5122 | 3.55 | 543.18758 |

| | | | | | | | |
|---|---|---|---|---|---|---|---|
| 65 | QL | 915.4567 | [M+H]+ | C45H70O19 | 914.4489 | 3.65 | 498.87668 |
| 66 | QL | 915.4581 | [M+H]+ | C45H70O19 | 914.4503 | 3.71 | 500.40521 |
| 67 | QL | 1386.6535 | [M+NH4]+ | C63H100O32 | 1368.6195 | 3.67 | 470.19728 |
| 68 | QL | 1076.4899 | [M+NH4]+ | C50H74O24 | 1058.4559 | 3.67 | 455.0902 |
| 69 | QL | 1062.5129 | [M+NH4]+ | C50H76O23 | 1044.4789 | 3.68 | 482.72355 |
| 70 | QL | 1048.5319 | [M+NH4]+ | C50H78O22 | 1030.4979 | 3.69 | 507.2807 |
| 71 | QL | 960.5157 | [M+NH4]+ | C47H74O19 | 942.4817 | 3.73 | 536.89908 |
| 72 | QL | 858.4452 | [M+NH4]+ | C42H64O17 | 840.4112 | 3.75 | 518.61202 |
| 73 | QL | 1224.6002 | [M+NH4]+ | C57H90O27 | 1206.5662 | 3.76 | 490.11914 |
| 74 | QL | 1092.5579 | [M+NH4]+ | C52H82O23 | 1074.5239 | 3.75 | 516.32356 |
| 75 | QL | 1192.5372 | [M+NH4]+ | C55H82O27 | 1174.5032 | 3.79 | 450.46813 |
| 76 | QL | 959.5209 | [M+H]+ | C48H78O19 | 958.5131 | 3.87 | 541.62515 |
| 77 | QL | 812.4423 | [M+NH4]+ | C41H62O15 | 794.4083 | 4.06 | 544.40789 |
| 78 | QL | 830.4894 | [M+NH4]+ | C42H68O15 | 812.4554 | 4.08 | 589.29109 |
| 79 | QL | 830.4901 | [M+NH4]+ | C42H68O15 | 812.4561 | 4.22 | 590.13346 |
| 80 | QL | 670.4518 | [M+NH4]+ | C36H60O10 | 652.4178 | 4.38 | 673.87395 |
| 81 | QL | 816.5099 | [M+NH4]+ | C42H70O14 | 798.4759 | 4.96 | 624.48722 |
| 82 | PG | 975.515 | [M+H]+ | C48H78O20 | 974.5072 | 3.66 | 527.92627 |
| 83 | PG | 827.442 | [M+H]+ | C42H66O16 | 826.4342 | 3.79 | 534.17641 |
| 84 | QL | 829.4568 | [M+H]+ | C42H68O16 | 828.449 | 3.65 | 550.7219 |
| 85 | QL | 974.495 | [M+NH4]+ | C47H72O20 | 956.461 | 3.83 | 507.9554 |
| 86 | QL | 1324.5802 | [M+NH4]+ | C60H90O31 | 1306.5462 | 3.79 | 438.02557 |

**Wilding MD**. The Relative Saponin Content of Alfalfa Varieties in Utah as Measured by Chick Growth Depression. : 46.
