## Appendix S2 for "A domestic plant differs from its wild relative along multiple axes of within-plant trait variability and diversity"

**Supplementary Information for**

**Title:** A domestic plant differs from its wild relative along multiple axes of within-plant trait variability and diversity

**Authors:** Moria L. Robinson[1,2,5]*, Anthony L. Schilmiller[3], William C. Wetzel[1,2,4,5,6]

[1]Department of Entomology, Michigan State University

[2]Kellogg Biological Station, Michigan State University

[3]Mass Spectrometry and Metabolomics Core, Michigan State University

[4]Department of Integrative Biology, Michigan State University

[5]Ecology, Evolution, and Behavior Program, Michigan State University

[6]AgBioResearch, Michigan State University

**This supplement includes:**

Supplementary text
Figures S2.1 to S2.2

**Table S2.1.** Structures of top models used to estimate differences in among-leaf trait variability and trait means between wild and domestic plants, among all leaves per plant (N=9). This table permits comparison of estimates (effect sizes) between domestication and covariates (trait mean; plant size; percent wild subspecies ancestry) when model selection included these variables in top models. See Table S2.2 for results within leaf age class (young, middle, old). For each trait, there are three models: variability (standard deviation; chemical diversity) predicted by domestication status alone (top); variability (standard deviation; chemical diversity) predicted by domestication status and the accompanying trait mean (or total compound concentration) (middle); and trait mean (or total compound concentration) predicted by domestication status (third row). NA indicates that a term was not relevant for a given global model; hyphens indicate terms that were included in model selection, but not included in the top model. Note that only two of the three wild subspecies were included as potential terms in models, as all three would force collinearity (see Table S1.4). Estimates and 95% confidence intervals (parentheses) are shown. For random effects, we show the intraclass correlation coefficient (ICC) for all random terms in the model. See Tables S1-S2 for main effects (wild vs. domestic) in terms of % change, and for interpretation of interactions. Significance levels are indicated by asterisks: $p \leq .05$ (*), $p \leq .01$ (**), and $p \leq .001$ (***). Note that some very small coefficients or CIs are shown as “0.00” due to truncation/rounding during table generation.

| | | | *fixed effects* | | | | | | | | *random effects* | | |
|---|---|---|---|---|---|---|---|---|---|---|---|---|---|
| **Trait** | **Basic model structure / Hypothesis** | **Intercept** | **Domestic /wild** | **Domestic/wild × Mean** | **Mean (or total compound concentration)** | **Plant size (volume)** | **Repro /veg** | **% ssp. *falcata*** | **% ssp. *caerulea*** | **Integration method** | **Random effect(s)** | **ICC** | **nObs** |
| SLA | SD ~ domestication | 3.58 (2.6 /4.5)*** | 3.01 (1.9/4.2)** | NA | NA | --- | --- | 0.01 (0.0/.03) | --- | NA | NA | NA | 60 |
| | SD ~ domestication + mean | -5.48 (-7.8/-3.1)*** | 1.71 (0.8/2.6)** | --- | 0.54 (0.4/0.7)** | --- | --- | 0.01 (0.0/.02) | --- | NA | NA | NA | 60 |
| | mean ~ domestication | 16.87 (15.8/17.9)*** | 2.29 (0.8/3.8)** | NA | NA | --- | --- | --- | --- | NA | NA | NA | 60 |
| Trichome density | SD ~ domestication | 3.86 (3.1/4/6)*** | 0.06 (-1.0/1.1) | NA | NA | --- | --- | --- | --- | NA | NA | NA | 60 |
| | SD ~ domestication + mean | 0.19 (-0.1/0.5) | -0.04 (-0.5/0.4) | 0.10 (-0.0/0.2) | 0.19 (0.1/0.2)*** | --- | --- | --- | --- | NA | NA | NA | 60 |
| | mean ~ domestication | 4.84 (3.9/5.78)*** | -1.33 (-2.7/0.0)* | NA | NA | --- | --- | --- | --- | NA | NA | NA | 60 |
| LWC | SD ~ domestication | 4.92 (4.5/5.4)*** | -0.04 (-0.7/0.6) | NA | NA | --- | --- | --- | --- | NA | NA | NA | 60 |
| | SD ~ domestication + mean | 15.31(9.4/21.3)*** | 0.35 (-0.3/1.0) | --- | -0.15 (-0.2/-0.1)*** | --- | --- | --- | --- | NA | NA | NA | 60 |

| | | | | | | | | | | | | | |
|---|---|---|---|---|---|---|---|---|---|---|---|---|---|
| | mean ~ domestication | 68.74 (67.5/70.0)*** | 2.61 (0.9/4.4)** | NA | NA | --- | --- | --- | --- | NA | NA | NA | 60 |
| Carbon(C) | SD ~ domestication | 0.96 (0.8/1/1)*** | 0.14 (-0.1/0.4) | NA | NA | --- | --- | 0.003 (0.0/0.01) | --- | NA | NA | NA | 43 |
| | SD ~ domestication + mean | 4.06 (-2.7/10.8) | 0.13 (-0.1/0.4) | --- | -0.1 (-0.2/.1) | --- | --- | 0.003 (0.0/0.01)* | --- | NA | NA | NA | 43 |
| | mean ~ domestication | 43.8 (43.4/44.0)*** | -0.21(-0.6/-0.2) | NA | NA | --- | --- | --- | --- | NA | NA | NA | 43 |
| Nitrogen (N) | SD ~ domestication | 0.81 (0.7/0.9)*** | 0.11 (-0.0/0.3) | NA | NA | -0.001 (-0.0/0.0) | --- | --- | --- | NA | NA | NA | 43 |
| | SD ~ domestication + mean | -0.01 (-0.4/0.4) | 0.06 (-0.1/0.2) | NA | 0.25 (0.1/0.4)*** | 0.001 (-0.0/0.0) | --- | --- | --- | NA | NA | NA | 43 |
| | mean ~ domestication | 3.16 (2.9/3.4)*** | 0.24 (-0.0/0.5) | NA | NA | --- | --- | 0.004 (0.0/0.0)* | --- | NA | NA | NA | 43 |
| C:N | SD ~ domestication | 3.05 (2.7/3.4)*** | 0.41 (-0.1/0.9) | NA | NA | --- | --- | --- | --- | NA | NA | NA | 43 |
| | SD ~ domestication + mean | 1.12 (-1.7/3.9) | -1.4 (-4.9/2.0) | 0.14 (-0.1/0.4) | 0.14 (-0.1/0.3) | --- | --- | --- | --- | NA | NA | NA | 43 |
| | mean ~ domestication | 14.67 (13.8/15.6)*** | -0.67 (-1.8/0.4) | NA | NA | --- | --- | -0.02 (-0.0/-0.0)* | --- | NA | NA | NA | 43 |
| Saponins (single) | SD ~ domestication | -3.14 (-3.5/-2.8)*** | 0.32 (0.1/0.5)** | NA | NA | 0.003 (-0.0/0.0) | --- | --- | --- | -0.51 (-1.0/-0.1)* | mean\|compound; mean\|plant | 0.41 | 2892 |
| | SD ~ domestication + mean | -2.84 (-3.0/-2.7)*** | -0.01 (-0.1/.1) | --- | 0.99 (0.9/1.0)*** | 0.001 (-0.0/0.0) | --- | --- | --- | -0.24 (-0.4/-0.1)*** | mean\|compound; mean\|plant | 0.06 | 2892 |
| | mean ~ domestication | -2.57 (-2.9/-2.3)*** | 0.37 (0.2/0.6)*** | NA | NA | 0.002 (-0.0/0.0) | --- | --- | --- | -0.35 (-0.76/0.07) | compound; plant | 0.40 | 2892 |
| Saponins (total) | SD ~ domestication | 4.87 (3.7/6.1)*** | 2.10 (0.6/3.6)** | NA | NA | --- | --- | -0.02 (-0.0/0.0) | --- | NA | NA | NA | 60 |
| | SD ~ domestication + mean | 1.95 (0.4/3.6)* | 0.82 (-0.5/2.2) | --- | 0.19 (0.1/0.3)*** | --- | --- | -0.01 (-0.0/0.0) | --- | NA | NA | NA | 60 |
| | mean ~ domestication | 15.92 (12.4/19.4)*** | 6.04 (1.4/10.6)* | NA | NA | 0.02 (-0.0/0.1) | --- | -0.03 (-0.1/0.0) | --- | NA | NA | NA | 60 |
| Richness | richness ~ domestication | 59.17 (56.3/62.1)*** | 3.91 (0.1/7.8)* | NA | NA | 0.02 (-0.0/0.1) | --- | -0.02 (-0.1/0.0) | --- | NA | NA | NA | 60 |
| | richness ~ domestication + total | 61.33 (59.0/63.6)*** | -0.69 (-3.9/2.5) | --- | 8.77 (6.2/11.3)*** | 0.01 (-0.0/0.1) | --- | -0.02 (-0.1/0.0) | --- | NA | NA | NA | 60 |
| Shannon diversity | Shannon ~ domestication | 2.80 (2.7/2.9)*** | -0.06 (-0.2/0.1) | NA | NA | 0.000 (-0.0/0.0) | --- | --- | -0.002 (-0.0/-0.0)* | NA | NA | NA | 60 |
| | Shannon ~ domestication + total | 2.80 (2.7/2.9)*** | -0.06 (-0.3/0.1) | --- | 0.004 (-0.0/-0.0) | 0.00 (-0.0/0.0) | --- | --- | 0.002 (-0.0/-0.0)* | NA | NA | NA | 60 |
| Simpson diversity | Simpson ~ domestication | 0.89 (0.8/0.9)*** | -0.03 (-0.1/0.0) | NA | NA | -0.00 (-0.0/0.0) | --- | --- | -0.0004 (-0.0/-0.0)* | NA | NA | NA | 60 |
| | Simpson ~ domestication + total | -0.89 (0.8/0.9)*** | -0.04 (-0.1/0.0) | --- | 0.00008 (-0.0/0.0) | -0.0004 (-0.0/0.0) | --- | --- | -0.0004 (-0.0/-0.0)* | NA | NA | NA | 60 |
| Beta (0/1) | Beta ~ domestication | 0.06 (0.0/0.1)*** | 0.002 (-0.0/0.0) | NA | NA | --- | --- | --- | --- | NA | NA | NA | 60 |
| | Beta ~ domestication + total | 0.08 (0.1/0.1)*** | 0.01 (-0.0/0.0) | NA | -0.001 (-0.0/-0.0)** | --- | --- | --- | --- | NA | NA | NA | 60 |
| Beat (prop) | Beta ~ domestication | -1.77 (-1.9/-1.7)*** | -0.15 (-0.3/-0.0)* | NA | NA | --- | --- | --- | -0.001 (-0.0/0.0) | NA | NA | NA | 60 |
| | Beta ~ domestication + total | -1.76 (-1.9/-1.6)*** | -0.15 (-0.3/-0.0)* | --- | -0.004 (-0.0/0.0) | --- | --- | --- | --- | NA | NA | NA | 60 |
| Beta (raw) | Beta ~ domestication | 0.22 (0.2/0.2)*** | -0.02 (-0.0/0.0) | NA | NA | --- | --- | --- | -0.0002 (-0.0/0.0) | NA | NA | NA | 60 |
| | Beta ~ domestication + total | 0.24 (0.2/0.3)*** | -0.004 (-0.0/0.0) | --- | -0.002 (-0.0/-0.0)** | --- | --- | --- | -0.0002 (-0.0/0.0) | NA | NA | NA | 60 |

**Table S2.2.** Structures of top models used to estimate differences in among-leaf trait variability and trait means between wild and domestic plants, within leaf age class (young, middle, old). This table permits comparison of estimates (effect sizes) between domestication and covariates (trait mean; plant size; percent wild subspecies ancestry) when model selection included these variables in top models. For each trait, there are three models: variability (standard deviation; chemical diversity) predicted by domestication status alone (top); variability (standard deviation; chemical diversity) predicted by domestication status and the accompanying trait mean (or total compound concentration) (middle); and trait mean (or total compound concentration) predicted by domestication status (third row). NA indicates that a term was not relevant for a given global model; hyphens indicate terms that were included in model selection, but not included in the top model. Note that only two of the three wild subspecies were included as potential terms in models, as all three would force collinearity (see Table S1.4). Estimates and 95% confidence intervals (parentheses) are shown. For random effects, we show the intraclass correlation coefficient (ICC) for all random terms in the model. See Tables S1-S2 for main effects (wild vs. domestic) in terms of % change, and for interpretation of interactions. Significance levels are indicated by asterisks: $p \leq .05$ (*), $p \leq .01$ (**), and $p \leq .001$ (***). Note that some very small coefficients or CIs are shown as "0.00" due to truncation/rounding during table generation.

| | | | *Fixed effects* | | | | | | | | | | *Random effects* | | |
|---|---|---|---|---|---|---|---|---|---|---|---|---|---|---|---|
| Trait | Basic model structure / Hypothesis | Intercept | Domestic /wild | Leaf stage | Mean | Domestic/wild × Leaf stage | Domestic/wild × Leaf stage × Mean | Plant size (volume) | Repro /veg | % ssp. *falcata* | % ssp. *caerulea* | Integration method | Random effect(s) | ICC | nObs |
| SLA | SD ~ domestication | 4.85 (4.1/5.6)*** | 1.44 (0.7/2.2)*** | -1.56 (-2.5/-0.7) [middle]** -3.04 (-3.94/-2.13) [old]*** | NA | --- | NA | --- | --- | --- | --- | NA | --- | --- | 180 |
| | SD ~ domestication + mean | -0.95 (-2.7/0.8) | 0.79 (0.1-1.5)* | -0.46 (-1.3/0.4) [middle] -1.21 (-2.2/-0.3)* [old] | 0.29 (0.2-0.4)** | --- | --- | --- | --- | --- | --- | NA | --- | --- | 180 |
| | mean ~ domestication | 18.6 (17.2/20.0)*** | 5.7 (3.7/7/7)*** | -1.49 (-3.19/0.22) [middle] -3.69 (-5.39/-1.99) [old]*** | NA | -4.73 (-7.1/-2.3) [dom×middle]*** -5.44 (-7.9/-3.0) [dom×old] | NA | --- | --- | --- | --- | NA | plant | 0.29 | 180 |
| Tr ic | SD ~ domestication | 0.97 (0.6/1.4)*** | -0.3 (-0.7/0.2) | -0.28 (-1.2/-0.4) [middle]*** -1.89 (-2.3/-1.5) [old]*** | NA | --- | NA | --- | --- | --- | --- | NA | --- | --- | 180 |

| | | | | | | | | | | | | | | | |
|---|---|---|---|---|---|---|---|---|---|---|---|---|---|---|---|
| | SD ~ domestication + mean | -0.82<br>(-1.1/-0.5)*** | 0.15<br>(-.1/-.3) | 0.21 (-.1/0.5) [middle]<br>0.31 (-.04/0.7) [old] | 0.87<br>(0.8/1.0)[log]<br>*** | --- | --- | --- | --- | --- | --- | NA | --- | --- | 180 |
| | mean ~ domestication | 8.92<br>(7.9/10.0)*** | -1.33<br>(-2.7/0.0)* | -5.27 (-6.1/-4.4) [middle]***<br>-6.99 (-7.8/-6.1) [old]*** | NA | --- | NA | --- | --- | --- | --- | NA | plant | 0.47 | 180 |
| LWC | SD ~ domestication | 4.75<br>(3.9/5.6)*** | -0.36<br>(-1.5/0.8) | -0.33 (-1.5/0.8) [middle]<br>-2.24 (-3.4/-1.1) [old]*** | NA | 1.01 (-0.6/2.6) [dom×middle]***<br>1.76 (0.2/3.4) [dom×old]* | NA | --- | --- | --- | --- | NA | --- | --- | 180 |
| | SD ~ domestication + mean | 13.74<br>(8.6/19.0)*** | 0.93<br>(0.3/1.6)** | 0.45 (-0.3/1.3) [middle]<br>-1.0 (-1.8/-0.2) [old]** | -0.14<br>(-0.2/-0.1)*** | --- | --- | --- | --- | --- | --- | NA | --- | --- | 180 |
| | mean ~ domestication | 65.78<br>(64.3/67.2)*** | 5.45<br>(3.4/7.5)*** | 3.80 (2.4/5.2) [middle]***<br>5.10 (3.7/6.5) [old]*** | NA | -3.67 (-5.6/-1.7) [dom×middle]***<br>-4.84 (-6.8/-2.9) [dom×old]*** | NA | --- | --- | --- | --- | NA | plant | 0.55 | 180 |
| Carbon (C) | SD ~ domestication | 0.60<br>(0.4/0.7)*** | 0.02<br>(-0.1/0.2) | -0.002 (-0.2/0.2) [middle]<br>0.07 (-0.1/0.2) [old] | NA | --- | NA | 0.002<br>(0.0/0.0)* | --- | --- | --- | NA | --- | --- | 151 |
| | SD ~ domestication + mean | 3.14<br>(0.1/6.2)* | -0.04<br>(-0.2/0.1) | -0.04 (-0.2/0.1) [middle]<br>-0.03 (-0.2/0.2) [old] | -0.06<br>(-0.1/0.0) | --- | --- | 0.002<br>(-0.0/0.0) | --- | 0.002<br>(0.0/0.0)* | --- | NA | --- | --- | 151 |
| | mean ~ domestication | 44.44<br>(44.1/44.8)*** | 0.18<br>(-0.3/0.7) | -0.44 (-0.8/-0.1) [middle]*<br>-1.59 (-2.0/-1.2) [old]*** | NA | -0.66 (-1.2/-0.1) [dom×middle]*<br>-0.36 (-0.9/0.2) [dom×old] | NA | --- | --- | --- | --- | NA | plant | 0.53 | 151 |
| Nitrogen (N) | SD ~ domestication | 0.72<br>(0.6/0.9)*** | 0.05<br>(-0.1/0.2) | 0.12 (-0.1/0.3) [middle]<br>-0.27 (-0.4/-0.1) [old]*** | NA | --- | NA | 0.001<br>(-0.0/0.0) | --- | --- | --- | NA | --- | --- | 151 |
| | SD ~ domestication + mean | 0.13<br>(-0.2/0.5) | 0.02<br>(-0.1/0.2) | 0.07 (-0.1/0.2) [middle]<br>-0.22 (-0.4/-0.1) [old]** | 0.18<br>(0.1/0.3)*** | --- | --- | 0.001<br>(-0.0/0.0) | --- | --- | --- | NA | --- | --- | 151 |
| | mean ~ domestication | 2.99<br>(2.7/3.3)*** | 0.69<br>(0.3/1.1)*** | 0.59 (0.3/0.9) [middle]***<br>0.07 (-0.2/0.4) [old] | NA | -0.68 (-1.1/-0.2) [dom×middle]**<br>-0.71 (-1.1/-0.3) [dom×old]** | NA | -0.003<br>(-0.0/0.0) | --- | 0.003<br>(0.0/0.0)* | --- | NA | plant | 0.28 | 151 |
| C:N | SD ~ domestication | 2.92<br>(2.3/3.6)*** | -0.04<br>(-1.0/0.9) | -0.34 (-1.2/0.6) [middle]<br>-0.89 (-1.8/-0.0) [old]* | NA | 1.18 (-0.1/2.4) [dom×middle]<br>0.13 (-1.1/1.4) [dom×old] | NA | --- | --- | --- | --- | NA | --- | --- | 151 |
| | SD ~ domestication + mean | 4.05<br>(-0.0/8.1)* | -3.78<br>(-8.5/0.9) | -2.14 (-7.2/2.9) [middle]<br>-4.65 (-10.6/-1.3) [old] | -0.07<br>(-0.3/0.2) | 0.29 (-5.75/6.3) [dom×middle]<br>4.86 (-2.9/12.7) [dom×old] | 0.27 (-0.0/0.6) [mean]<br>0.13 (-0.2/0.5) [middle×mean]<br>0.26 (-0.1/0.7) [old×mean]<br>0.07 (-0.4/0.5) [dom×middle×mean]<br>-0.34 (-0.9/0.2) [dom×old×mean] | --- | --- | --- | --- | NA | --- | --- | 151 |
| | mean ~ domestication | 15.67<br>(14.6/16.8)*** | -2.07<br>(-3.5/-0.6)** | -2.50 (-3.7/-1.3) [middle]***<br>-1.30 (-2.5/-0.1) [old]* | NA | 2.64 (1.0/4.3) [dom×middle]**<br>2.35 (0.7/4.0) [dom×old]** | NA | --- | --- | -0.02<br>(-0.0/-0.0)* | --- | NA | plant | 0.27 | 151 |
| Saponins (single) | SD ~ domestication | -2.85<br>(-3.2/-2.5)*** | 0.42<br>(0.2/0.6)*** | -0.03 (-0.1/0.1) [middle]<br>0.14 (0.1/0.2) [old]** | NA | -0.07 (-0.2/0.1) [dom×middle]<br>0.06 (-0.1/0.2) [dom×old] | NA | --- | --- | -0.0003<br>(-0.0/0.0) | --- | -0.33<br>(-0.8/0.1) | mean\|compound;<br>mean\|plant | 0.38 | 8939 |
| | SD ~ domestication + mean | -3.63<br>(-3.7/-3.5)*** | -0.05<br>(-0.2/.1) | 0.05 (-0.01/0.1) [middle]<br>0.06 (0.0/0.1) [old]* | 1.01<br>(0.98/1.04)*** | -0.08 (-0.2/-0.0) [dom×middle]*<br>-0.10 (-0.2/-0.0) [dom×old]** | --- | --- | --- | -0.002<br>(-0.0/-0.0)** | --- | -0.24<br>(-0.4/-0.1)*** | mean\|compound;<br>mean\|plant | 0.20 | 8939 |
| | mean ~ domestication | -2.86<br>(-3.2/-2.5)*** | 0.42<br>(0.3/0.6)*** | -0.06 (-0.1/0.0) [middle]<br>0.17 (0.1/0.2) [old]*** | NA | --- | NA | --- | --- | --- | --- | -0.33<br>(-0.8/0.1) | compound; plant | 0.38 | 8939 |
| Saponins (total) | SD ~ domestication | 1.10<br>(0.8/1.4)*** | 0.66<br>(0.2/1.1)** | -0.30 (-0.7/0.1) [middle]<br>-0.17 (-0.5/0.2) [old] | NA | -0.45 (-1.0/0.0) [dom×middle]<br>-0.41 (-0.9/0.1) [dom×old] | NA | 0.003<br>(-0.0/0.0) | --- | -0.005<br>(-0.0/-0.0)** | NA | NA | plant | 0.15 | 180 |
| | SD ~ domestication + mean | 0.54<br>(0.2/0.9)*** | 0.13<br>(-0.1/0.4) | -0.35 (-0.6/-0.1) [middle]**<br>-0.36 (-0.6/-0.1) [old]** | 0.04<br>(0.0/0.1)*** | --- | --- | 0.002<br>(-0.0/0.0) | --- | -0.004<br>(-0.0/-0.0)* | NA | NA | plant | 0.09 | 180 |
| | mean ~ domestication | 2.53<br>(2.3/2.8)*** | 0.57<br>(0.3/0.9)*** | -0.12 (-0.3/0.0) [middle]<br>0.04 (-0.1/0.2) [old] | NA | -0.13 (-0.3/0.1) [dom×middle]<br>0.01 (-0.2/0.2) [dom×old] | NA | 0.0007<br>(-0.0/0.0) | --- | -0.0008<br>(-0.0/0.0) | NA | NA | plant | 0.77 | 180 |
| Richness | richness ~ domestication | 50.0<br>(48.0/52.0)*** | 4.18<br>(2.6/5.7)*** | -0.73 (-2.6/1.1) [middle]<br>-1.53 (-3.4/0.3) [old] | NA | --- | NA | --- | --- | -0.03<br>(-0.1/-0.0)* | -0.02<br>(-0.1/-0.0)* | NA | NA | NA | 540 |
| | richness ~ domestication + total | 49.8<br>(47.3/52.4)*** | -0.21<br>(-3.6/3.2) | 0.10 (-1.4/1.6) [middle]<br>-1.10 (-3.1/0.8) [old] | 6.58<br>(5.1/8.1)*** | 0.91 (-0.7/2.5) [dom×middle]<br>-1.2 (-2.9/0.5) [dom×old] | --- | --- | --- | --- | --- | NA | mean\|leaf stage<br>mean\|plant | 0.79 | 540 |
| Shannon diversity | Shannon ~ domestication | 2.62<br>(2.6/2.7)*** | -0.04<br>(-0.1/0.0) | 0.10 (0.0/0.2) [middle]**<br>0.03 (-0.0/0.1) [old] | NA | --- | NA | 0.00<br>(-0.0/0.0) | --- | --- | -0.002<br>(-0.0/-0.0)*** | NA | NA | NA | 540 |
| | Shannon ~ domestication + total | 2.59<br>(2.5/2.7)*** | -0.05<br>(-0.2/0.1) | 0.02 (-0.0/0.1) [middle]<br>0.02 (-0.0/0.1) [old] | -0.05<br>(-0.1/0.0) | 0.15 (0.1/0.2) [dom×middle]***<br>0.05 (-0.0/0.1) [dom×old] | 0.08 (-0.0/0.2) [mean]<br>-0.1 (-0.2/-0.0) [middle×mean]**<br>-0.14 (-0.2/-0.1) [old×mean]***<br>-0.00 (-0.1/0.1) [dom×middle×mean]<br>0.04 (-0.1/0.2) [dom×old×mean] | --- | --- | --- | -0.002<br>(-0.0/0.0) | NA | mean\|plant | 0.79 | 540 |

| | | | | | | | | | | | | | | | |
|---|---|---|---|---|---|---|---|---|---|---|---|---|---|---|---|
| Simpson diversity | Simpson ~ domestication | 0.84<br>(0.8/0.9)*** | -0.02<br>(-0.0/-0.0)*** | 0.02 (0.0/0.0) [middle]**<br>-0.001 (-0.0/0.1) [old] | NA | --- | NA | --- | 0.02<br>(0.0/0.0)*<br>[veg] | 0.0002<br>(-0.0/0.0)* | -0.0003<br>(-0.0/-0.0)** | NA | NA | NA | 540 |
| | Simpson ~ domestication + total | 0.84<br>(0.8/0.9)*** | -0.05<br>(-0.1/-0.0)* | -0.001 (-0.0/0.0) [middle]<br>-0.01 (-0.0/0.0) [old] | -0.01<br>(-0.0/0.0) | 0.04 (0.0/0.1) [dom×middle]***<br>0.02 (0.0/0.1) [dom×old]* | 0.02 (-0.0/0.1) [mean]<br>-0.02 (-0.0/-0.0) [middle×mean]*<br>-0.03 (-0.1/-0.0) [old×mean]***<br>-0.01 (-0.0/0.0) [dom×middle×mean]<br>0.00 (-0.0/0.0) [dom×old×mean] | --- | --- | 0.0003<br>(-0.0/0.0) | --- | NA | mean\|plant | 0.73 | 540 |
| Beta (0/1) | Beta ~ domestication | -3.60<br>(-3.8/-3.4)*** | 0.26<br>(-0.0/0.6) | 0.18 (-0.0/0.4) [middle]<br>0.28 (0.1/0.5) [old]** | NA | -0.44 (-0.7/-0.2) [dom×middle]<br>-0.48 (-0.8/-0.2) [dom×old]** | NA | --- | --- | --- | --- | NA | plant | 0.56 | 180 |
| | Beta ~ domestication + total | -3.41<br>(-3.7/-3.1)*** | 0.36<br>(0.1/0.7)* | 0.14 (-0.1/0.4) [middle]<br>0.27 (0.1/0.5) [old]** | -0.01<br>(-0.0/-0.0)* | -0.47 (-0.8/-0.2) [dom×middle]***<br>-0.46 (-0.8/-0.2) [dom×old]** | NA | --- | --- | --- | --- | NA | plant | 0.51 | 180 |
| Beta (prop) | Beta ~ domestication | -2.52<br>(-2.7/-2.4)*** | -0.22<br>(-0.4/-0.1)** | 0.1 (-0.0/0.2) [middle]<br>0.05 (-0.1/0.2) [old] | NA | --- | NA | --- | --- | --- | --- | NA | plant | 0.11 | 180 |
| | Beta ~ domestication + total | -2.43<br>(-2.6/-2.3)*** | -0.18<br>(-0.3/0.0)** | 0.08 (-0.1/0.2) [middle]<br>0.05 (-0.1/0.2) [old] | -0.01<br>(-0.0/0.0) | --- | --- | --- | --- | --- | --- | NA | plant | 0.07 | 180 |
| Beta (raw) | Beta ~ domestication | -2.26<br>(-2.4/-2.1)*** | 0.10<br>(-0.1/0.3) | 0.03 (-0.2/0.2) [middle]<br>0.09 (-0.1/0.3) [old] | NA | -0.20 (-0.5/0.1) [dom×middle]<br>-0.37 (-0.7/-0.1) [dom×old]** | NA | --- | --- | --- | --- | NA | plant | 0.16 | 180 |
| | Beta ~ domestication + total | -1.98<br>(-2.2/-1.8)*** | 0.14<br>(-0.1/0.4) | -0.004 (-0.2/0.2) [middle]<br>0.08 (-0.1/0.3) [old] | -0.01<br>(-0.0/-0.0)** | -0.23 (-0.5/0.1) [dom×middle]<br>-0.36 (-0.7/-0.1) [dom×old]* | --- | --- | --- | -0.003<br>(-0.0/-0.0)*** | --- | NA | plant | 0.01 | 180 |
